# Atomic scale description of the allosteric coupling between a lipid bilayer and a membrane protein

**DOI:** 10.1101/2025.01.24.634742

**Authors:** Clarisse Fourel, Yanna Gautier, Alexandre Pozza, François Giraud, Elodie Point, Christel Le Bon, Karine Moncoq, Guillaume Stirnemann, Jérôme Hénin, Ewen Lescop, Laurent J. Catoire

## Abstract

Biological membranes are complex environments whose functions are closely tied to the dynamic interactions between lipids and proteins. Here, we utilize high-pressure NMR of lipid nanodiscs paired with molecular dynamics simulations to elucidate at the atomic scale the allosteric dialog between the lipid bilayer and a bacterial model membrane protein, OmpX. We discover that OmpX delays the gelation process by liquefying the annular shell of lipids through hydrophobic and roughness matching processes at the protein surface. Furthermore, the alteration of the mechanical properties of the lipid bilayer directly impacts the energy landscape of amino acid side chains at the lipid/protein interface, but also, unexpectedly, at the protein core. Our work highlights a potential thermodynamically coupled but kinetically uncoupled allosteric pathway linking lipid dynamics with the interior of membrane proteins, directly impacting our understanding of membrane function.

**Graphical Abstract:** 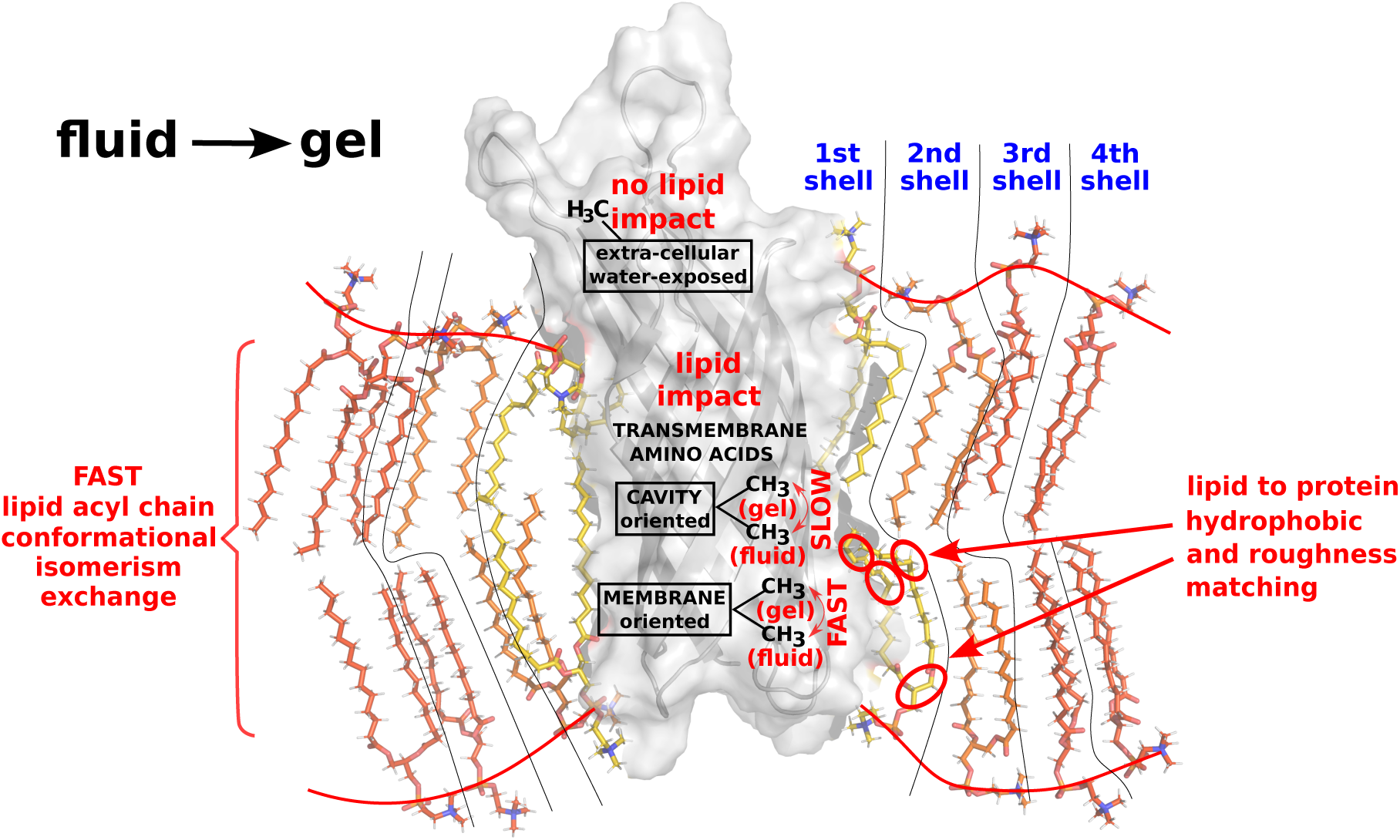

## Introduction

Cell and organelle membranes are intricate homeostatic regulatory systems. They are mainly composed of a diverse array of lipids, which interact with a wide variety of integral and peripheral proteins across different cell types, subcompartments of membrane organelles, and metabolic states^1^. A fundamental area of research focuses on the regulation of integral membrane proteins (IMPs) by the lipid bilayer^2^. Until now, most atomistic studies of IMP/lipid interactions have focused on high-affinity IMP/lipid contacts thanks to an ever increasing number of high-resolution 3D structures. However, how bulk membrane properties influence IMP free energy conformational landscape at the atomic scale is still poorly understood. In turn, how does the protein perturb the lipid behavior and response to pressure (P)/temperature (T) thermodynamic conditionsProgress in this area is currently limited by the difficulty of concomitantly characterizing lipids and proteins with high time and spatial resolutions, all on an atomic scale.

We propose a combination of three main approaches to address these questions. The first involves using soluble nano-objects (nanodiscs), where the protein is embedded within a lipid bilayer. Combining this with cutting-edge IMP isotope labeling and lipids at natural isotope abundance enables solution-state nuclear magnetic resonance (NMR) spectroscopy to reveal atomic details on both lipids and IMP. The second key aspect is the use of hydrostatic pressure to control the dynamics of the lipid phase^3–5^, including fluid-gel phase transitions, to probe potential correlated motion between proteins and lipids^6^. The third cornerstone of our approach is the atomic-level interpretation of the experimental measurements using all-atom molecular dynamics simulations, which have become indispensable tools in a wide range of biophysical applications.

By combining these three essential components, we investigate the complex relationships between one integral membrane protein and lipids at both the molecular and atomic levels. We focus on the collective properties of the annular lipid shell surrounding the protein and its long-distance impact on the IMP conformational energy landscape. We provide a clear rationalization at the atomic and molecular levels of how membrane proteins influence the behavior of lipids, their response to pressure, and the spatial extent of this perturbation. In turn, our strategy highlights the reciprocal impact of lipid phase transitions on membrane protein structure and dynamics. Unequivocally, our study reveals an allosteric coupling between bilayer mechanics and subtle conformational changes in the membrane protein. These physical changes affect not only the amino acid side chains in direct contact with the lipids, but also those facing the protein core, away from the lipids. Our detailed description of the interplay between lipid phase transitions and IMPs at the atomic/molecular scale aims to advance our understanding of the allosteric effects of lipids on membrane protein function.

## Results and Discussion

All experimental and simulation data presented in this study comes from high-resolution NMR spectra of IMP-free or IMP-containing 1,2-dimyristoyl-sn-glycero-3-phosphocholine (DMPC) or 1,2-dimyristoleoyl-sn-glycero-3-phosphocholine (Δ9–cis–PC) MSP1D1 nanodiscs. DMPC and its unsaturated analog Δ9–cis–PC have very similar structures, except for one additional unsaturation in each acyl chain of Δ9–cis–PC (Fig. 1a). This results in a large decrease in gel/fluid transition temperature for Δ9–cis–PC (*T_m_* < 0°C) compared to DMPC (*T_m_*≃ 25°C) at 1 bar (Supplementary Fig. 1). Simulated data was obtained by explicit all-atom simulations of periodic DMPC or Δ9–cis–PC bilayers, with or without IMP. For this study, we chose the Outer membrane protein X (OmpX) from *Escherichia coli* as the model for IMP, as that protein has been extensively characterized by NMR spectroscopy, including in DMPC nanodiscs of equivalent dimensions, at ambient^7^ and high pressure^6^. OmpX is perdeuterated and contains ^13^CH_3_ specifically incorporated in Ala/Val/Ile residues allowing us to obtain high quality 1D ^1^H and 2D ^1^H-^13^C NMR correlation spectra, which simultaneously revealed OmpX and lipid (DMPC or Δ9–cis–PC) behavior (see Supplementary Fig. 2 and 3).

**Fig. 1.**
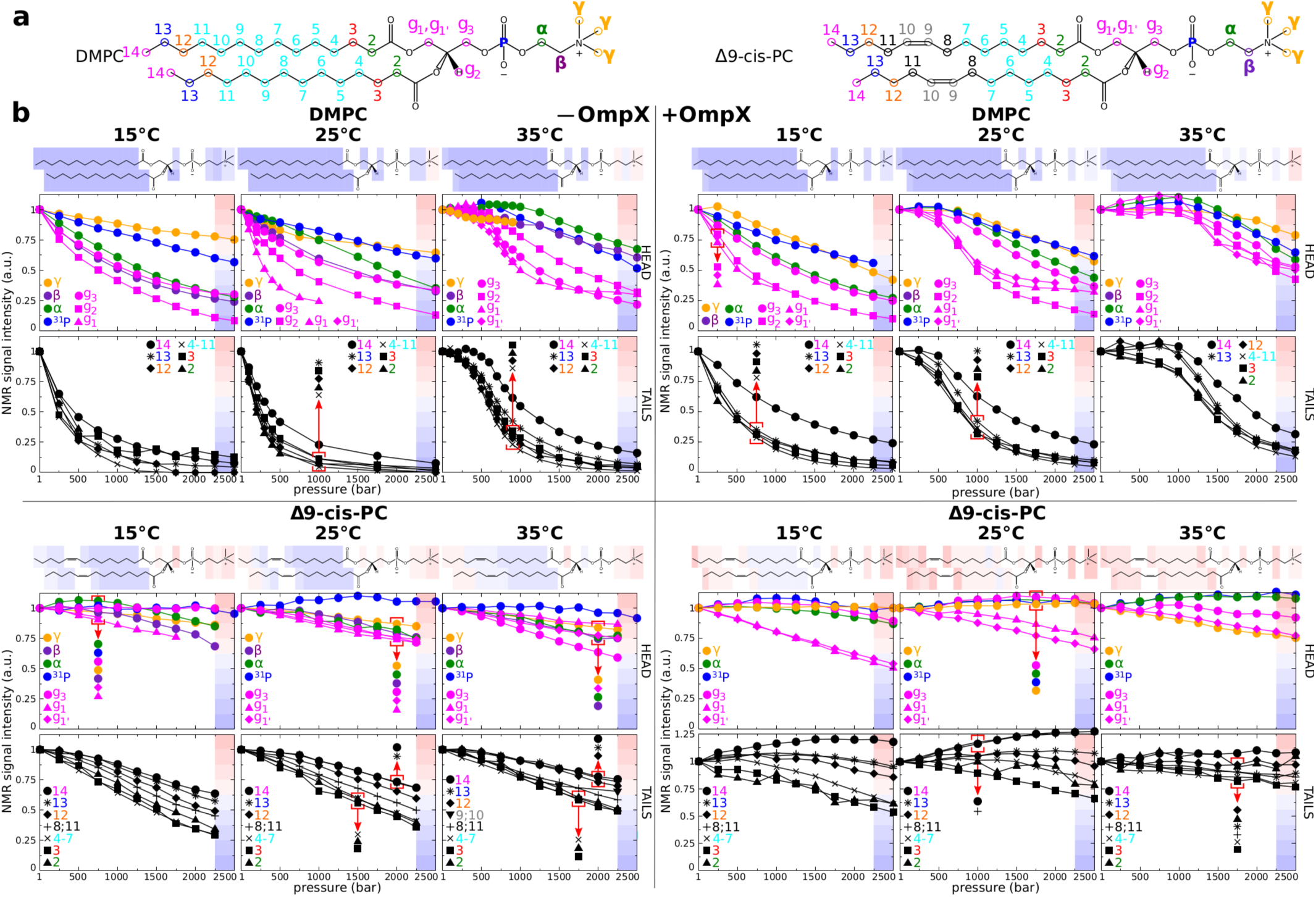
Barotropic behavior of DMPC and Δ9–cis–PC ^1^H NMR signal intensities in nanodiscs in the absence or presence of the protein OmpX. **a,** Primary chemical structures of DMPC and Δ9–cis–PC with atom nomenclature used in this work. b, Barotropic evolutions of NMR signal intensities at 15, 25, and 35°C observed for DMPC (upper left and right panels) and Δ9–cis–PC (lower left and right panels) nanodiscs in the absence (upper and lower left panels) or presence (upper and lower right panels) of OmpX. Above each graph is represented a lipid chemical structure overlaid with colored rectangles. The color code corresponds to the signal intensity range per 0.125 of NMR signal intensity observed at the end of the pressure ramp. By convention, all intensities are scaled to 1 at ambient pressure for easy comparison.

### DMPC and Δ9–cis–PC bilayer phase behavior

We previously demonstrated^6^ that ^1^H NMR can detect the fluid-to-gel transition upon pressurization of DMPC in nanodiscs at 40°C. Here, we expanded our exploration of the fluid-gel phase boundary of DMPC in presence or absence of OmpX using extended temperature (15-35°C) and pressure (1-2500 bar) ranges (Fig. 1b). In the absence of OmpX, a clear phase transition is observed at 35°C (near ∼600 bar) but not at 25°C and 15°C. This is in full agreement with the DMPC phase diagram defined in infinite bilayers^8^ (Supplementary Fig. 1) and previous DSC studies on DMPC nanodiscs ^9^. The presence of OmpX makes a clear phase transition visible around 1250 bar at 35°C and around 600 bar at 25°C. No transition was observed at 15°C. This clearly shows that (at least part of) the DMPC lipids surrounding OmpX undergo a cooperative fluid-to-gel transition but with a phase boundary globally shifted by ∼650 bar when compared with OmpX-devoid nanodiscs. Furthermore, negative controls using Δ9–cis–PC demonstrate no discernible transitions in the presence or absence of OmpX (Fig. 1). Collectively, this strongly suggests that Δ9–cis–PC remains fluid at all P/T values, both in the presence or absence of OmpX.

We gained atomic details on the lipid phase transition and the impact of the insertion of OmpX by carrying out all-atoms MD simulations. These simulations revealed a pressure-induced phase transition consistent with NMR data. We compared low and high pressure DMPC systems both with and without an embedded OmpX molecule (Fig. 2). At 25°C, the fluid-to-gel transition in OmpX-devoid DMPC is evident through the sharp decrease in area per lipid (from 60 to 50 Å^2^), *gauche* fraction (from ∼28% to ∼14%) and glycerol head hydration number when pressure increased from 1 to 2500 bar (Fig. 2b), which are all hallmarks of lipid gelation^10^. The transition is even sharper at 35°C and occurs between 500 and 750 bar which perfectly matches the transition observed experimentally by NMR (*P_m_* ≃ 600 bar). As a negative control, Δ9–cis–PC does not undergo any transition in simulations.

**Fig. 2.**
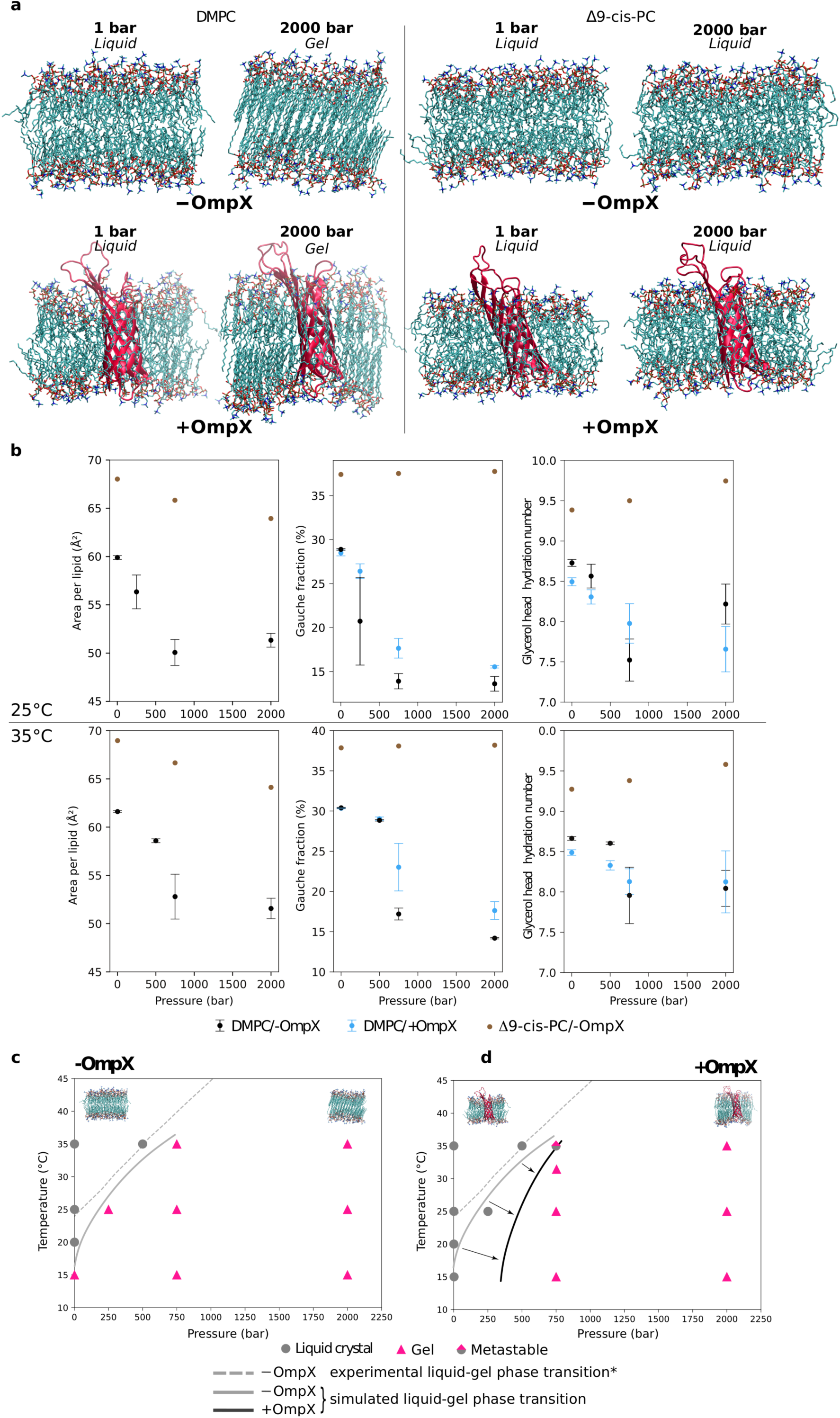
Pressure-induced phase transition of DMPC in the presence and absence of OmpX, characterized by simulations. **a,** Molecular rendering of simulated systems: pure DMPC (*left*) and Δ9-cis-PC (*right*) bilayers in the absence (top) or presence (bottom) OmpX (cartoon representation in red). In each case, snapshots in thermodynamic conditions corresponding to a liquid state and to a gel phase are shown. **b,** Area per lipid, *gauche* fraction and glycerol hydration number over the last 300 ns of the simulation as a function of pressure, at a temperature of 25°C and 35°C (data at 15°C is shown in Extended Data Fig. 1), averaged over three replicas (error bars: standard deviation). **c,d,** DMPC phase diagram obtained from the MD simulations (**c**) without and (**d**) with OmpX, with symbols corresponding to the liquid, gel, or metastable states of the system in the simulations. *corresponds to the experimental DMPC liquid-gel phase transition from Ichimori et al^8^.

### Effect of protein on lipid phase transition: dynamic plasticity of hydrophobic and roughness matching

MD simulations clearly demonstrate that OmpX impacts the lipid dynamics and phase transition of DMPC, preventing the decrease in *gauche* fraction at high pressures. Thus, OmpX favors the fluid phase, as observed experimentally (Fig. 2b). The simulated P/T phase diagram in absence (Fig. 2c) or presence of OmpX (Fig. 2d – full data in Extended Data Fig. 1) is fully consistent with the experimentally determined ones (Fig. 1b and ref^8^). In the Δ9–cis–PC control system, the *gauche* fraction is almost constant (36%), which is consistent with the total absence of gelation of Δ9–cis–PC around OmpX, as observed by NMR (Fig. 1b).

MD simulations also provide high spatial resolution of protein-induced perturbations of the lipids (Fig. 3). Regardless of the P/T conditions, the first two lipid solvation shells around the protein remain essentially liquid as judged from their specific *gauche* fraction values (Fig. 3a). Under all P/T conditions except low temperature (15°C) and pressure (1 bar), lipids beyond the second shell show *gauche* fraction values typical of those of a gel state, showing that the spatial extent of OmpX influence on lipid dynamics depends on P/T thermodynamics conditions, but also includes the first two solvation shells. Finally, for systems that have gelified in the presence of OmpX, the *gauche* fraction of lipids beyond the second solvation shell is very similar to that of the pure DMPC system. Simulations thus provide evidence that the roughness of protein surface cannot easily accommodate lipids in the extended all-*trans* conformations for the first two solvation shells.

**Fig. 3.**
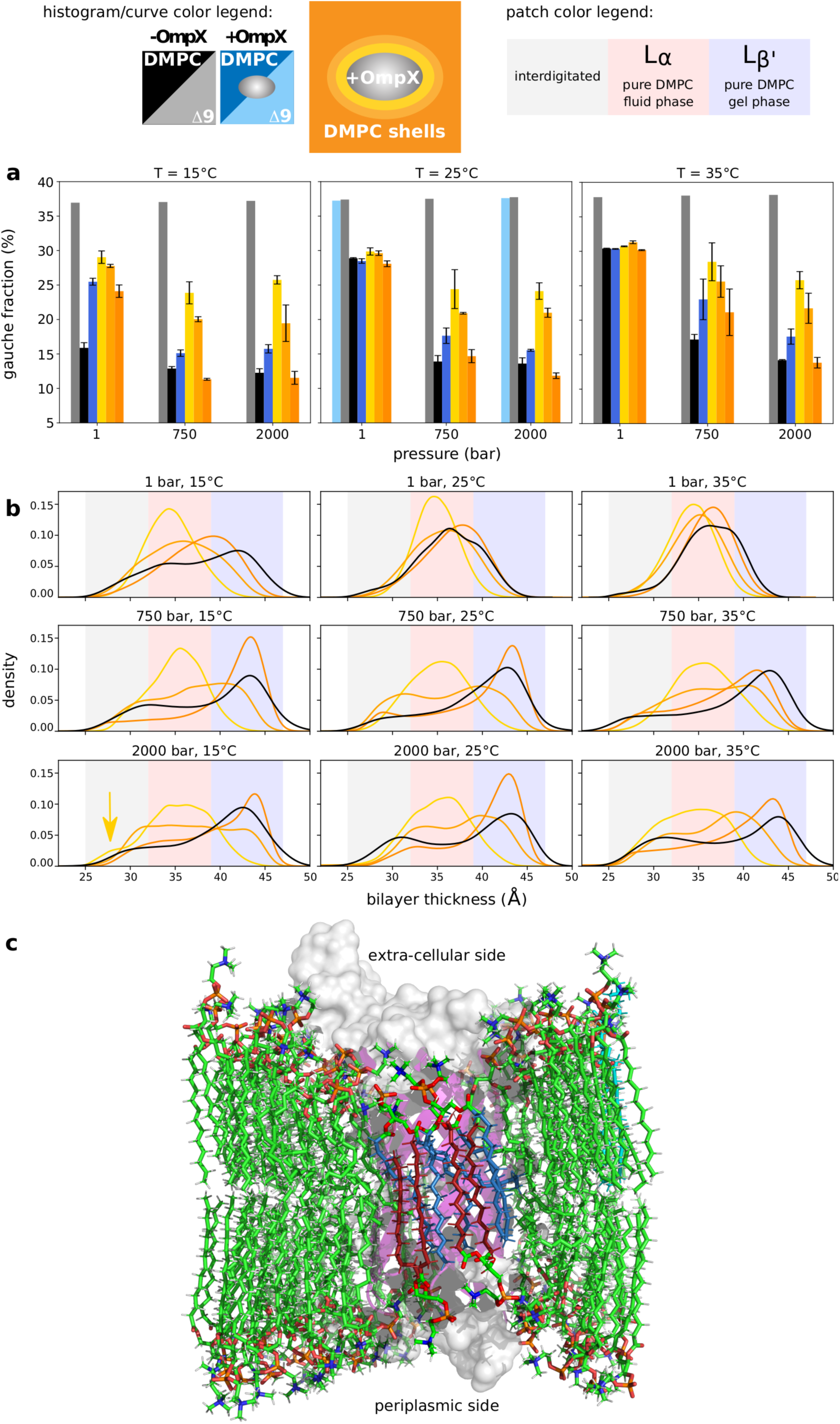
Distance-dependent perturbation of the lipid phase transition by OmpX. a) *gauche* fraction in acyl chains of the pure DMPC (black) or Δ9–cis–PC (gray) bilayers; in the entire DMPC bilayer with OmpX (dark blue) or Δ9–cis–PC with OmpX (light blue); and in solvation shells of OmpX in lipids (first shell: yellow; second shell: light orange; beyond the second shell: dark orange). Each panel shows plots for increasing pressures at a given temperature. Data points are the average among three replicas for DMPC (a single replica was simulated for Δ9–cis–PC), with error bars corresponding to the SD among these replicas. b) Bilayer thickness distributions as a function of temperature and pressure. Color codes are the same as above. Note that liquid bilayers (e.g., at 35°C/1 bar) exhibit one broad peak around 35 Å, whereas gel phases (e.g., at 15°C/2000 bar) are described by a bimodal distribution, with a main peak around 42 Å for the textbook L*_β_′* phase, and a secondary peak at low thickness (≃ 30 Å) corresponding to interdigitated regions. These spatial distributions were obtained by interpolating the bilayer boundaries on a regular grid, and sampling their spacing (see Methods). c) Molecular rendering of the organization of a gel-phase (15°C, 2 kbar) DMPC bilayer around OmpX. The protein surface is colored in gray and the the cartoon representation of the *β*-barrel in purple. To help to visualize interdigitation of lipid acyl chains and the match between the hydrophobic thickness of the bilayer and that of the protein (low thickness region indicated by the arrow in panel b), some acyl chains of the lipids of the first lipid shell were colored in blue (outer leaflet) and red (inner leaflet). According to Luzzati’s nomenclature^11^ L*_α_* designs a fluid-like crystalline phase that has a lamellar structure and conformationally disordered acyl chains in contrast with the L*β* gel phase that features acyl chains in a more extended conformation.

Bilayer thickness also strongly depends on the lipid phase. The bilayer is thicker in the gel phase and thinner in the fluid phase. Consequently, the fluid-to-gel transition for the DMPC-only bilayer is reflected by the increase in thickness at 15°C/2000 bar (gel phase) compared to 35°C/1 bar (fluid phase) (see Fig. 3b). In the presence of OmpX, the first two shells are liquid-like (i.e., thinner bilayer), while the third shell is more like the lipid distribution without OmpX. A clear distinction emerges between the first and second shells, particularly under high-pressure and low-temperature conditions, where the second shell displays a broader distribution, akin to that of a gel phase. Two additional observations can be made. First, the first shell is thinner than the corresponding liquid phase in the absence of OmpX, suggesting that lipids adapt to accommodate the presence of the protein. Second, the gel phase exhibits a bimodal and broad thickness distribution, with a primary peak corresponding to the well-ordered gel phase at large bilayer thickness and a significant population at much smaller thickness values, which can be attributed to interdigitation of the two lipid shells.

We can thus identify two types of perturbation induced by OmpX in DMPC bilayers. The first type is a short-range effect on the first two solvation shells that always remain disordered. This may originate from the roughness of the OmpX transmembrane surface, which would destabilize the extended conformers of the surrounding lipids and induce structural defects; and/or from the necessary hydrophobic matching between the lipid bilayer and the protein transmembrane surface, which would disrupt the organization of the lipids (Fig. 3c). The second type of protein-induced perturbation extends beyond these two shells and shifts the phase transition to higher pressures.

### Correlated structural and dynamic changes in OmpX with lipid phase transition

We also monitored OmpX ^1^H and ^13^C NMR signals from the same samples and NMR experiments as previously described^6^. This offered a unique opportunity to assess potential protein conformational changes concomitant to DMPC fluid-gel transition. Thanks to the *β*-barrel geometry of OmpX, we were able to distinguish and compare NMR signals of methyl groups in close contact with the lipid bilayer (residues V5, V39, I65, I73, I79, V83, V144, see Fig. 4), those pointing towards the interior of the cavity (A10, I40, V82, I141, see Fig. 5-6), and those located at the extracellular edge of the *β*-barrel (I132, V137). We aim to determine potential changes that correlate with the DMPC lipid phase transition visible at 25°C and at 35°C, using Δ9–cis–PC as a negative control. V135, which is located at the apex of the 8^th^ strand and exposed to water, serves as a reliable reference because the barotropic evolution of its NMR signal is very similar in DMPC and Δ9-cis-PC bilayers, showing that V135 is essentially not sensitive to membrane dynamic changes (Supplementary Fig. 4).

**Fig. 4.**
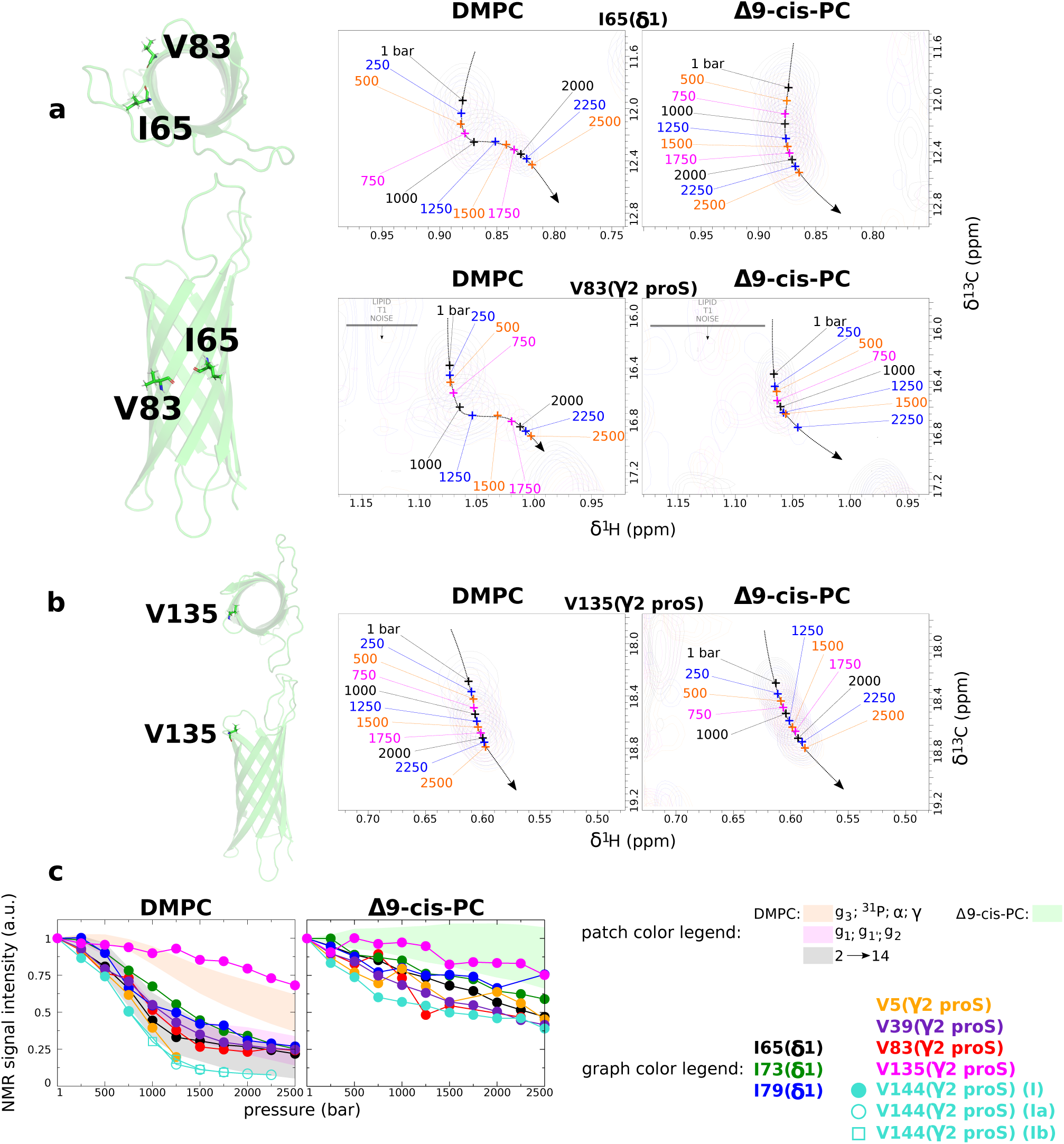
Barotropic evolution of ^1^H NMR signals of membrane-oriented ^13^CH_3_ of OmpX in DMPC and Δ9–cis–PC nanodiscs at 25°C. **a**, Illustrations of superimposed 2D ^1^H,^13^C SOFAST-HMQC NMR spectra^12^ spectra for residues I65 and V83 (see the same illustrations for V5, V39, I73, I79, V144 in Supplementary Fig. 5. For V144, see also Supplementary Fig. 10). The numbers represent the hydrostatic pressures that were applied. All spectra are represented on the same intensity scale. The positions of the amino acids are denoted in cartoon representations of OmpX, observed from both a parallel and a perpendicular axis (from the extracellular side) to the plane of the membrane. **b**, Same as **a** for the reference water-exposed residue V135. **c**, Comparison of the barotropic evolutions of ^13^CH_3_ NMR signal intensity for the membrane-oriented methyls groups and the reference water-exposed V135 either in DMPC (left) and Δ9–cis–PC (right) at 25°C. The colored patches depict the envelope of the barotropic evolution of intensity for different lipid NMR signals (from Fig. 1b).

**Fig. 5.**
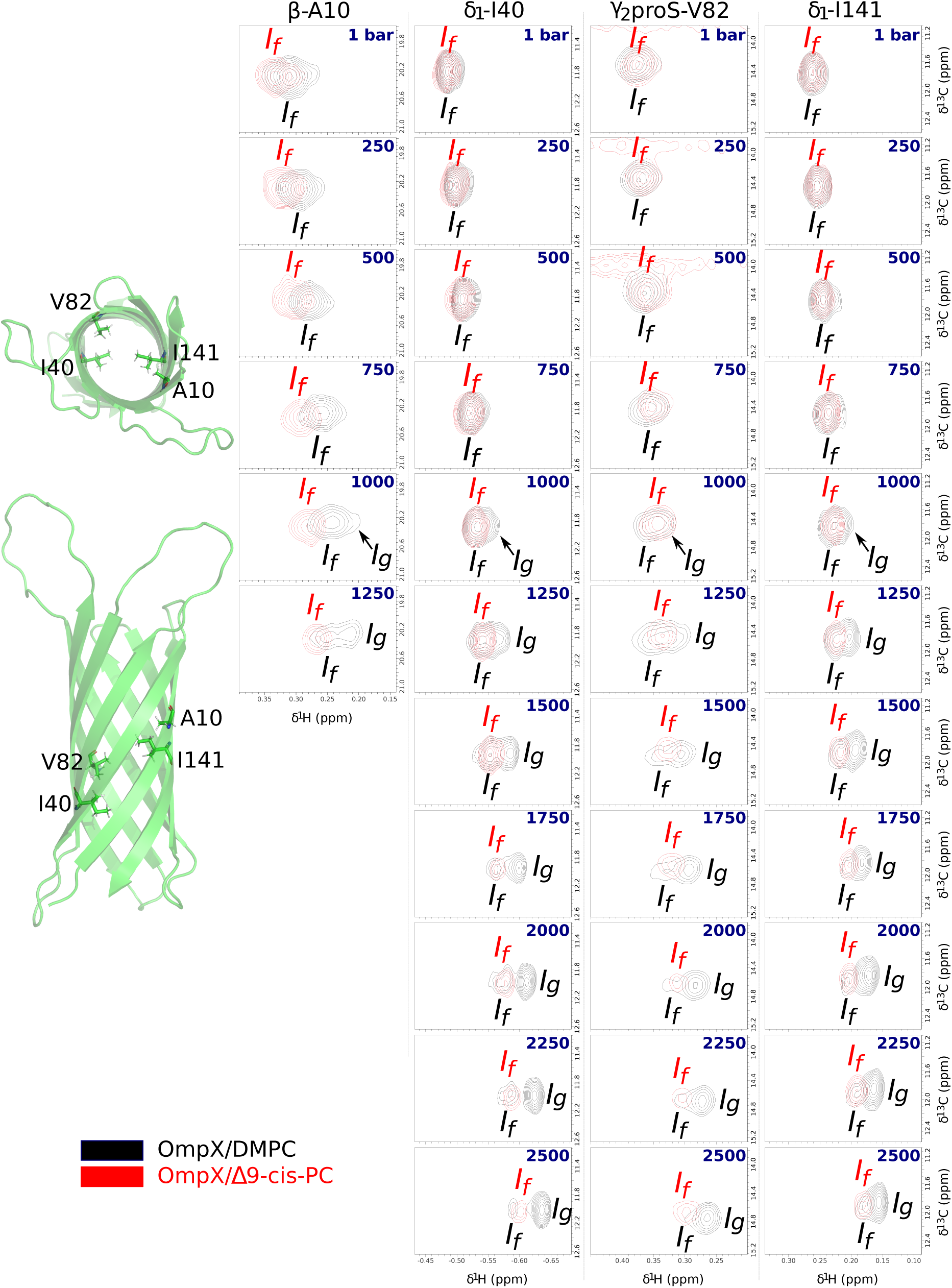
Barotropic evolution of ^1^H NMR signals of barrel interior-oriented ^13^CH_3_ of OmpX in DMPC (in black) and Δ9–cis–PC (in red) nanodiscs at 25°C (see also Supplementary Fig. 12). All the spectra are represented at the same scale in both dimensions (i.e., 0.25 and 1.5 ppm in respectively the ^1^H and ^13^C dimensions). *I_f_* and *I_g_* design the protein states when DMPC is in the fluid and gel states, respectively. The numbers in blue correspond to the hydrostatic pressure applied to the NMR sample. On the left are shown a top view (from the extracellular side) and a side view (from an axis parallel to the lipid bilayer) of a cartoon of OmpX (PDB ID: 2m06^7^) with the four cavity oriented residues represented in sticks.

**Fig. 6.**
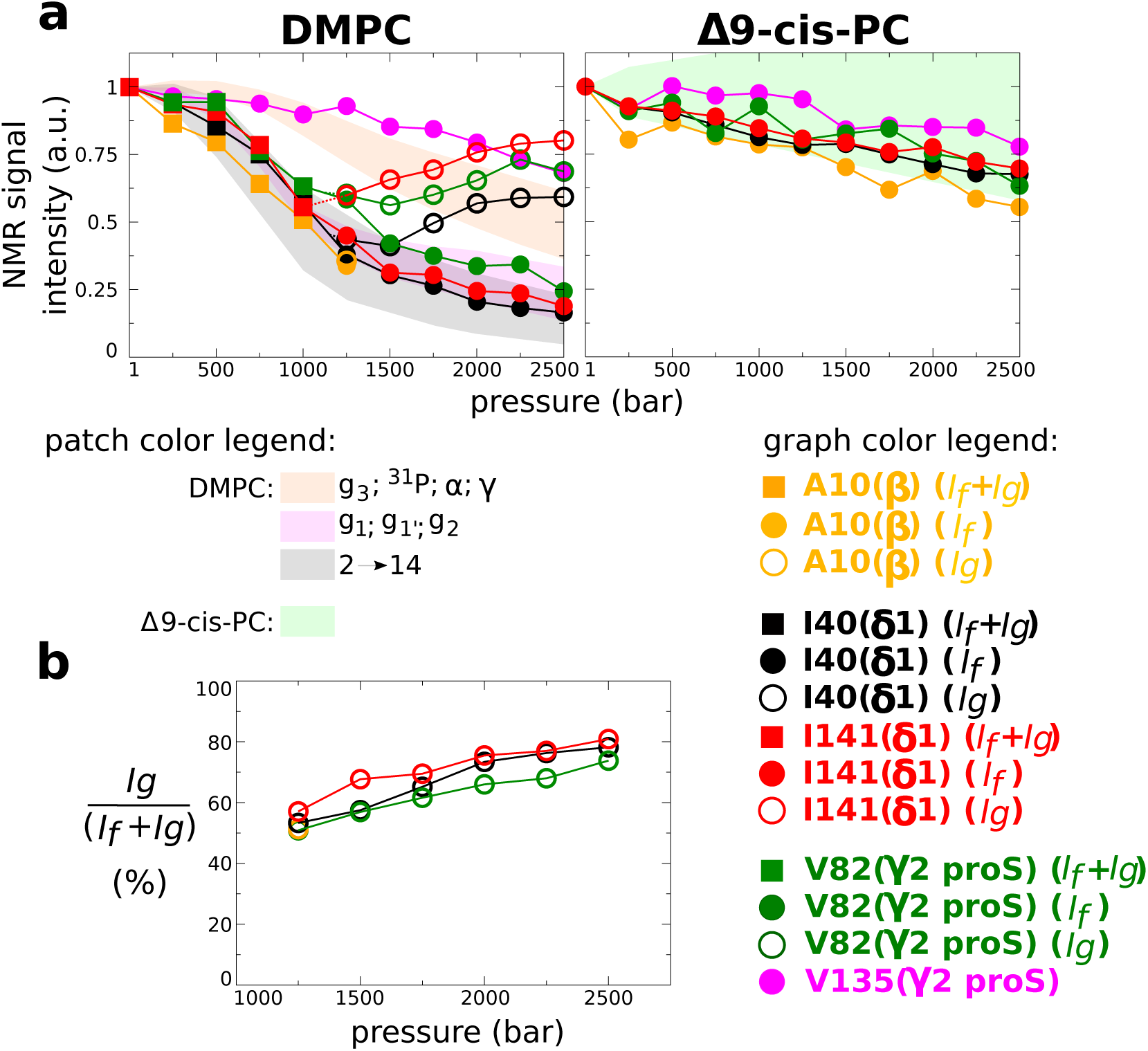
Barotropic evolution of NMR signal intensities of barrel interior-oriented ^13^CH_3_ of OmpX in DMPC and Δ9–cis–PC nanodiscs at 25°C (see also Supplementary Fig. 12). **a**, Comparison of the barotropic evolutions of ^13^CH_3_ NMR signal intensity. **b**, Barotropic evolution of the relative population of *I_g_* .

Fig. 4a depicts the barotropic progression, at 25°C, of the NMR signals of the well-resolved membrane- oriented I65-*δ*_1_ and V83-*γ*_2_proS methyls in DMPC and Δ9–cis–PC nanodiscs (I73, I79, V5, V39 and V144 NMR signals are displayed in Supplementary Fig. 5). Confirming and expanding our previous study^6^, we show here that DMPC gelation is associated with a significant decrease in intensity for the methyl groups that are directly exposed to the membrane. This decrease in methyl signal intensity is very similar to that of DMPC acyl chains (Fig. 4c). In contrast, this decrease in intensity for I65-*δ*_1_ and V83-*γ*_2_proS methyls is much reduced in Δ9–cis–PC lipids. This unequivocally shows that the relaxation change of these membrane-oriented methyl groups is directly connected to the changes in dynamics of the neighboring DMPC acyl chains upon phase transition and is not directly due to the applied pressure. The pressure-induced line broadening is most likely due to the gradual constriction of void volumes within this system.

We then analyzed methyl ^1^H and ^13^C chemical shift evolution. Chemical shifts generally report on the local chemical environment including the local protein structure or the organization of water/lipid solvation shells. Fig. 4a and Supplementary Fig. 5 clearly show that these evolutions display a clear sigmoidal inflection in DMPC whereas peaks shift almost linearly with pressure in Δ9–cis–PC. In DMPC, the sigmoidal inflection precisely corresponds to the evolution of DMPC acyl chain during DMPC phase transition. Additional experiments at 15 (Supplementary Fig. 6) and 35°C (Supplementary Fig. 7), and/or using the control Δ9–cis–PC nanodiscs further unambiguously established the coupling between the lipid phase transition in DMPC with both the dynamics and the chemical environment of membrane-oriented methyl groups.

Fig. 5 and 6 are the counterparts of Fig. 4 for amino acid side chains pointing towards the interior of the *β*–barrel. Remarkably, the barotropic evolution of the NMR signal of all well-resolved peaks (A10, I40, V82, I141) is also strongly sensitive to the DMPC main phase transition, despite the absence of direct contact with the lipids. Indeed, an additional NMR signal (*I_g_*) emerges gradually after the midpoint (around 750-1000 bar) of the lipid phase transition at 25°C (Fig. 5), while the barotropic evolution of the intensity of the initial signal observed at ambient pressure matches those corresponding to acyl chain protons of DMPC (Fig. 6a). At pressures higher than 2000 bar, *I_g_* is the predominant state (Fig. 5 and 6b). These observations strongly suggest that the methyl groups inside the cavity of the *β*-barrel feel, at least partially, the state of the lipid phase in concert with *I_f_* and *I_g_* corresponding to OmpX experiencing essentially a fluid and a gel phase, respectively. The emergence of *I_g_* at a higher pressure (∼1250bar) when temperature is increased to 35°C is further evidence of this concerted lipid and protein behavior (Extended Data Fig. 2 and Supplementary Fig. 8). As expected, negative controls with Δ9–cis–PC showed no signal splitting (Fig. 5) confirming that pressure alone is not sufficient to induce the emergence of *I_g_*signals. Comparison of methyl peak intensities at 1 and 2500 bar (Fig. 6a) in DMPC suggests that the major factor governing peak intensity changes upon gelation of DMPC is *I_f_* →*I_g_* population transfer, related to DMPC fluid→gel process, and not to changes in dynamics / relaxation due to gelation (which is the dominant process for membrane-oriented methyls). The similarity of methyl ^13^C chemical shifts for *I_f_* and *I_g_* implies a similar *χ*_2_ dihedral angle^13, 14^, so the observed Δ*δ*^1^H is due to the structural rearrangement of either the same methyl group, e.g. with a different *χ*_1_ dihedral angle and/or of the amino acids located in the vicinity of these methyl groups within OmpX interior. The OmpX crystal structure^15^ shows that the side chains of the three residues I40, V82, I141 can adopt at least two different rotamers in the cavity (Supplementary Fig. 9) allowing them to adjust their structure to changes in the physical properties of the bilayer. The very similar relative population changes with pressure (Fig. 6b) and the structural proximity for I40, I141 and V82 further strongly suggest a concerted conformational change along the *β*-barrel.

Our NMR analysis definitively demonstrates the allosteric protein conformational change upon DMPC gelation induced either by high pressure or low temperature. We observed only one NMR signal for each lipid atom, suggesting fast exchange at the ^1^H chemical shift regime (faster than millisecond timescale) between all chemical environments explored by the lipids, including the different phases and contacts with lipoprotein and IMP. In contrast, the exchange between *I_f_*and *I_g_*states appears to be at the slow exchange timescale (slower than millisecond timescale), suggesting thermodynamically (partially) coupled but kinetically uncoupled processes between lipid phase behavior and protein conformational change in the cavity. The uncoupled kinetics most likely reflect an allosteric pathway connecting lipids and OmpX core involving intermediate states connected through high energy barriers. This is in accordance with our MD simulations as attempts to employ enhanced sampling MD techniques along the relevant dihedral degrees of freedom to understand the impact of the lipid phase transition on protein side-chain motions. These failed to converge reliably across multiple replicas, highlighting that the conformational landscape of the side chains is influenced by the relaxation of slow (unknown) degrees of freedom in the surrounding environment. Addressing this challenge more accurately will require further methodological developments and investigations.

V144 *γ*_2_-proS methyl, which is located at the periplasmic edge of the 8^th^ *β*-strand, is exposed to the bilayer based on our MD simulations. It also explores an additional conformation *I_a_*, at the slow timescale, when lipids are in the gel phase (Supplementary Fig. 10). This strongly suggests that V144 dynamics is connected kinetically to the conformational change with the protein core. Isoleucines I132 and V137 are spatially close to each other at the two extremities of the extracellular *β*_7_-*β*_8_ strands at the edge of the bilayer and show an even more complex behavior. These methyls explore at 25°C two distinct conformations *I* and *II* at the slow chemical shift timescale with state *II* being disfavored at high pressure, irrespective of the lipid phase. On top of this, state *I*, but not state *II*, further splits into two additional states *I_a_* and *I_b_*when DMPC transitions to the gel, with *I_a_*being the predominant state at higher pressures (Supplementary Fig. 11). This reveals a complex conformational landscape, partially dependent on membrane dynamics, at the *β*_7_-*β*_8_/loop L4 boundary that contains several residues related to bacterial virulence^15^ (Supplementary Fig. 2).

## Conclusion

Membranes of living cells are typically crowded by integral and peripheral membrane proteins segregating in areas of various lipid and/or protein compositions. Each area has distinct membrane physical properties such as fluidity or thickness, and functions. In our study, we show that by introducing a defect in the bilayer organization, OmpX locally perturbs lipids over ∼2 lipid shells. This effect is observed in both experimental and simulated data, and it increases the fluidity of neighboring lipids. This is explained by the tendency of lipids to adapt to the protein surface roughness and the matching of bilayer thickness to the IMP hydrophobic surface.

Our work provides quantitative evidence of the allosteric coupling of lipid dynamics with subtle conformational changes along the transmembrane region of the protein, including parts inside, i.e., away from the lipid bilayer. Given the well-known impact of membrane composition in IMP function^2^, this result thus highlights a potential mechanism by which lipid dynamics may allosterically control IMP function by fine-tuning conformational changes at their binding or active sites, even in the case of extremely rigid IMPs, such as OmpX. This approach also opens up a new way of studying a wide variety of fields such as mechanically activated membrane proteins^16, 17^, the adaptation of biological membranes to pressure in deep-sea organisms^18, 19^, and drug design^20^.

## Methods

### Production and purification of recombinant OmpX and MSP1D1 proteins

Bacterial expression in *E. coli* (BL21(DE3) strain) and purification of OmpX and the lipoprotein MSP1D1 were carried out as previously described in ^15^ and ^21^, respectively. For solution-state NMR experiments, OmpX was uniformly deuterated and ^15^N-labeled in M9 minimal media in 100%-^2^H_2_O (^2^H*>*99%, Eurisotop) solutions supplemented with 2 g/L of u-[^2^H,^12^C]D-glucose (^2^H≃98%, Eurisotop) as the source of carbon and 1 g/L of ^15^NH_4_Cl (^15^N≃98%, EURISO-TOP) as the nitrogen source. Specific labeling with ^13^C and protonated methyls at Ala, Val (*γ*2pro*S*) and Ile (*δ*_1_-Ile) residues was achieved by following a published protocol^22, 23^ (TLAM kit from NMRBio).

### NMR sample preparation

The nanodiscs were formed by adding the lipoprotein to detergent-solubilized DMPC or Δ9–cis–PC phospholipids in the absence or presence of OmpX. The formation of nanodiscs was achieved by trapping the detergent with polystyrene beads (Bio-Beads SM-2 Adsorbent Media, Bio-Rad)^7, 21^. The nanodiscs utilized in this study (based on MSP1D1 lipoprotein) have a diameter of approximately 10 nm which corresponds to a bilayer area of 4400 Å^2 21^. Such nanodiscs consist of ∼90 molecules of DMPC per leaflet. In the presence of OmpX, which has an ellipsoidal cross-sectional area of ∼600–700 Å^2^ in the transmembrane region based on the NMR structure in DMPC nanodiscs (PDB ID: 2M06^7^), each leaflet contains around 80 molecules of DMPC, thus surrounding the protein by 4 lipid shells.

The NMR buffer solution was 25 mM Tris-HCl (pH 7.4), 50 mM NaCl, 2 mM EDTA in 90% H_2_O/10% D_2_O. OmpX-free and OmpX-containing lipid disc concentrations were comprised between 300 and ∼500 *µ*M for a total volume of 300 *µ*L in a alumina-toughened zirconia NMR tube (Daedalus Innovations LLC).

### NMR spectroscopy

Solution-state NMR experiments were performed at 15, 25 and 35°C between 1 and 2500 bar. They were conducted on Avance III HD Bruker spectrometers operating at ^1^H 700 and 950 MHz, equipped with TXO and TCI cryoprobes, respectively. 1D ^1^H experiments were collected using the *zgesgp* Topspin pulseprogram. 2D ^1^H,^13^C SOFAST-HMQC^12^ spectra were acquired at 950 (with OmpX-containing DMPC or Δ9–cis–PC nanodiscs and with OmpX-free DMPC nanodiscs) and 700 MHz (with OmpX-free Δ9–cis–PC nanodiscs) with a 200 ms recycling delay, 100 ms acquisition time (*t*_2_*_max_*) in the direct dimension, and 13.4 ms (*t*_1_*_max_*) in the indirect dimension (data size = 256(*t*_1_)×2,456(*t*_2_) complex points). The variable flip angle for the PC9 shape pulse was set to 120°. The number of acquisitions per increment was 64, for a total experiment time of 1h22 min. All ^31^*P* experiments were conducted at 600 MHz with a TBI probe. Data processing and analysis were performed with TopSpin NMR software.

Pressure was adjusted by an Xtreme-60 Syringe Pump apparatus (Daedalus Innovations LLC). A layer of mineral oils was used as a barrier between the sample and the pressurizing medium as described in^24^. Before NMR data collection at high pressure, a rapid pressure jump to 2500 bar was done for each sample outside the spectrometer to ensure the absence of leakage. The pressure ramp was from 1 to 2500 bar, each 100 or 250 bar. A delay of 15 min was imposed after each pressure change and prior to NMR data collection to allow system equilibration. Kinetic experiments confirmed that the lipid system was equilibrated within this timeframe and one-dimensional ^1^H spectra collected before and after 2D data collection were identical indicating equilibrium was indeed reached. A complete pressure cycle typically corresponds to ∼24 hours of data collection.

### Simulated systems

Systems were built using the CHARMM-GUI Membrane Builder tools^25–28^. Membranes were modelled as infinite bilayer systems. All systems were then constructed using the same protocol: a rectangular box was built with the length of Z based on a water thickness of 22.5 Å and the length of X and Y equal to 60 Å.

### Molecular dynamics simulations

All MD simulations were performed using NAMD 3 version *α*7^29^ with the Charmm36m all-atom force field^30^ and the TIP3P water model. Simulations were carried out under periodic boundary conditions based on rectangular boxes containing a hydrated lipid bilayer (in the xy plane, i.e., normal to the z-axis) embedded or not with OmpX.

We used the WYF parameter for cation-*π* interactions^31^ and hydrogen mass repartitioning^32^ which allows the use of a 4 fs time step. The different systems (pure lipid bilayers and bilayers with OmpX) were first minimised and relaxed in the NPT ensemble (i.e. with a constant number of particles N, constant pressure P and constant temperature T), at 1 bar and 30.15°C. The files provided by the CHARMM-GUI Membrane were used for minimisation, consisting of 6 cycles of 90 ns each, with planar and dihedral restraints that are progressively removed over the cycles. The systems were then additionally relaxed with a time step of 4 fs for 400 ns for the systems with protein and for 10 ns for the pure lipid systems. The following parameters were used for the production run: a time step of 4 fs was used to integrate Newton’s equations of motion and all bond lengths were constrained. The simulations were performed in the NPT ensemble and a combination of pressures (1 bar, 250 bar, 500 bar, 750 bar and 2000 bar) and temperatures (15°C, 25°C and 35°C) were used to explore the DMPC phase diagram. Langevin thermostat and NAMD Nosé-Hoover Langevin piston^33, 34^ were used to control the temperature and pressure, respectively. Particle mesh Ewald were set with a grid spacing of 1.0. Non-bonding interactions were limited to 12.0 Å and smoothing functions were applied beyond 10 Å for both electrostatics and van der Waals forces. Pure lipid systems were simulated for 1 *µ*s and lipid bilayers with protein were simulated for 1.5 *µ*s, with 3 replicas of each system. This gives a total simulation time of 36 *µ*s for DMPC bilayers and 54 *µ*s for OmpX in DMPC. The control lipid was simulated for 9×1 *µ*s for the pure lipid bilayer and 2×1 *µ*s for the system including the protein. Configurations were saved every 100 ps for analysis.

### Trajectory analysis

Trajectories were analysed using VMD^35^ version 1.9.4 through Tcl scripts. Analyses were performed on the last 300 ns of each simulation, taking into account the time needed for the lipid phase transition to occur.

The box volume was normalised to the condition with the largest volume: 1 bar 35°C. The area per lipid is defined as the area of a leaflet (corresponding to the area of the computational box in the xy plane) divided by the number of lipids it contains. Dihedral angles of the lipid acyl chains were used to calculate the *gauche* fraction, which corresponds to the number of *gauche* angles (grouping the *gauche*+ (0-120°) and *gauche*− (240-360°) angles^36^) divided by the total number of dihedral angles in the system (11 dihedrals for each lipid chain for all lipids). The *gauche* fraction was used as an indicator of lipid phase transition. To observe the spatial organisation of the lipids around the protein we extracted the number of lipids within 15 Å of the protein at intervals of 0.1 Å over the last 500 ns of simulation. The central glycerol carbon was used as a reference for the lipid position and any beta-sheet heavy atom of the protein was taken to represent the protein. We calculated the atomic radial pair distribution function *g*(*r*) and the number integral 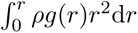 between the central glycerol carbon and each water oxygen atom to quantify the hydration of the lipid head. To do this we used the VMD ’measure gofr’ function on the last 300 ns of the simulation with the following parameters: delta=0.1, rmax=15, usepbc=1, first=6999, last=-1 and step=1. The area per lipid, as well as the volume of the simulation box, and as the *gauche* fraction were expressed as a function of time with running averages calculated using a convolution method with a window of 100 ns. The averaged presented in this paper were calculated on the data of each replica and the standard deviations were calculated between the replicas.

The effect of the protein on the local membrane environment was assessed by measuring the bilayer thickness. The entire simulation trajectory was centered, aligned and fitted without changing membrane orientation using the *α* carbons of the structured *β*-sheet portions of the protein. Lipids were marked as belonging to the “upper” or “lower” leaflet according to their position respectively to the bilayer midplane. The x, y and z coordinates of each phosphorus atom were then extracted for each frame of the simulation. For each leaflet, a 100×100 grid was constructed based on the x, y positions of the phosphorus atoms. Z positions of the phosphorus atoms were extrapolated by linear interpolation to create a plane corresponding to the surface of the upper and lower leaflet. The instantaneous distance between the upper and lower planes was then calculated and considered as the bilayer thickness. When the protein is inserted into the lipid bilayer, the center of mass of the upper (z-position atoms > 0) and lower (z-position atoms < 0) portions of the protein were extracted and a patch was created in the plane to mimic the position of the protein in each leaflet. To account for the different layers of lipids characterized around the protein, additional patches were created with radii of 10 Å and 18 Å for the first layer and 18 Å and 25 Å for the second layer. The bilayer thickness was measured every 0.1 ns for the last 300 ns of the simulation. Movies of the bilayer thickness were made showing the thickness calculated every nanosecond over the entire length of the simulation (1 microsecond for the pure lipid systems and 1.5 microseconds for the simulations of OmpX in the DMPC bilayer).

## Supporting information

Supplementary Informtion

## Data availability

Data supporting the findings of this work are available within the Article, Extended Data, Supplementary Information and source data files.

## Acknowledgements

We thank Daniel Picot for his helpful expertize in analyzing the electron density of OmpX with COOT software and Sebastien Billès for his contribution in the assignment of NMR lipid signals. This work was funded by the Centre National de la Recherche Scientifique (CNRS), Université Paris-Cité, the Agence Nationale de la Recherche (ANR-17-CE11-0011 & ANR-22-CE29-0020), Laboratoire d’Excellence (LabEx) DYNAMO (ANR-11-LABX-0011) and Equipements d’Excellence (EQUIPEX) CACSICE (ANR- 11-EQPX-0008) from the French Ministry of Research. Financial support from the IR INFRANALYTICS FR2054 CNRS for access to NMR spectrometers is gratefully acknowledged. This work was also supported by the French Infrastructure for Integrated Structural Biology (FRISBI) ANR-10-INBS-0005.

## Author contributions

G.S., J.H., E.L., L.J.C. conceived of the project. A.P., E.P., C.L. and K.M. performed biochemistry; A.P. and L.J.C. performed the NMR sample preparation. C.F., A.P., F.G, E.L. and L.J.C. performed the NMR data collection and analysis. Y.G., J.H. and G.S performed and analyzed the MD simulations. All authors discussed the data and their interpretation and edited the paper. The funders had no role in study design, data collection and analysis, decision to publish or preparation of the paper.

## Additional information

Supplementary Information –as a pdf file– and two Extended Data Figures are available for this manuscript.

---

**Extended Data Fig. 1.**
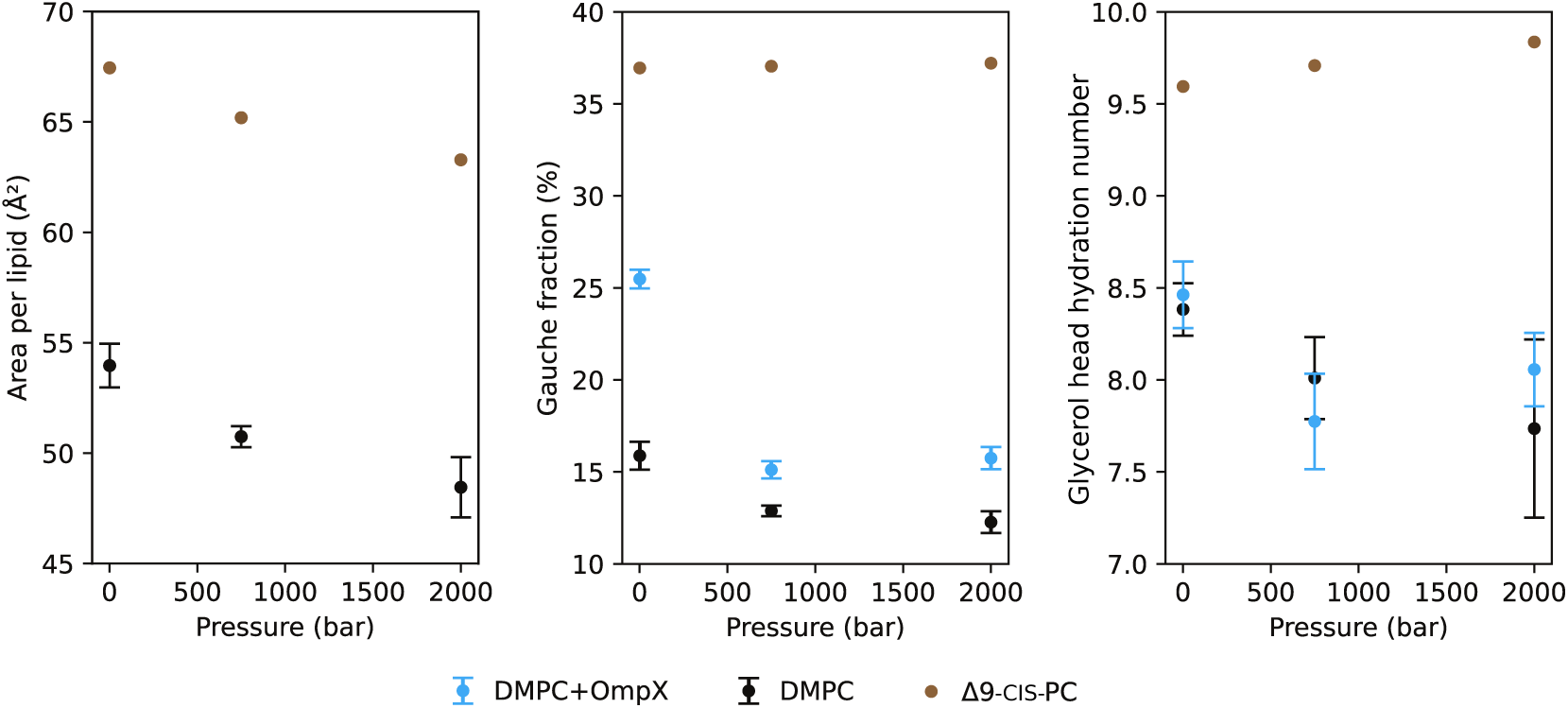
Additional simulation parameters details at 15°C (in complement to data at 25 and 35°C displayed in. Fig. 2b**). a**, Area per lipid for DMPC and Δ9–cis–PC bilayers, **b**, *gauche* fraction and **c**, Glycerol hydration number without and with OmpX in DMPC and Δ9–cis–PC bilayers. Data is calculated over the last 300 ns of the simulation as a function of pressure, averaged over three replicas (error bars: standard deviation).

**Extended Data Fig. 2.**
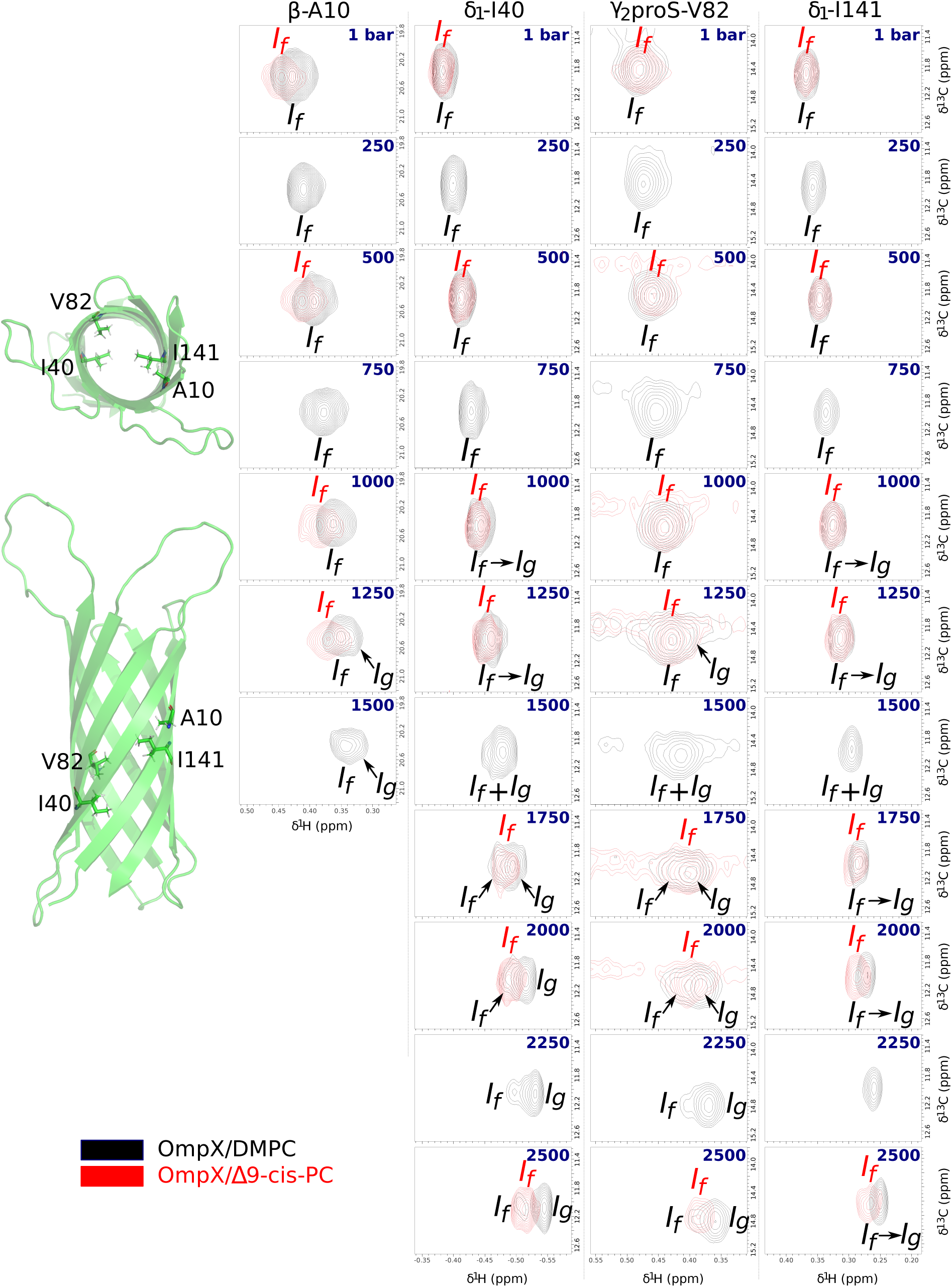
Barotropic evolution of NMR signals of barrel interior-oriented. ^13^**CH**_3_ **of OmpX in DMPC (in black) and** Δ**9–cis–PC (in red) nanodiscs at 35°C.** Legend same as Fig.5. Compared to data at 25°C (Fig. 5), I141 displays a faster chemical exchange between *I_f_* and *I_g_*. NMR experiments at 250, 750, 1500 and 2250 were not performed with OmpX/Δ9–cis–PC nanodiscs.

