## Supplementary Informtion for "Atomic scale description of the allosteric coupling between a lipid bilayer and a membrane protein"

---

#### Contents

1 Additional Figures 2

2 Analysis of pressure-induced evolution of OmpX $^{13}CH_3$ chemical shifts: definition of the angle $\theta$ 17

### 1 Additional Figures

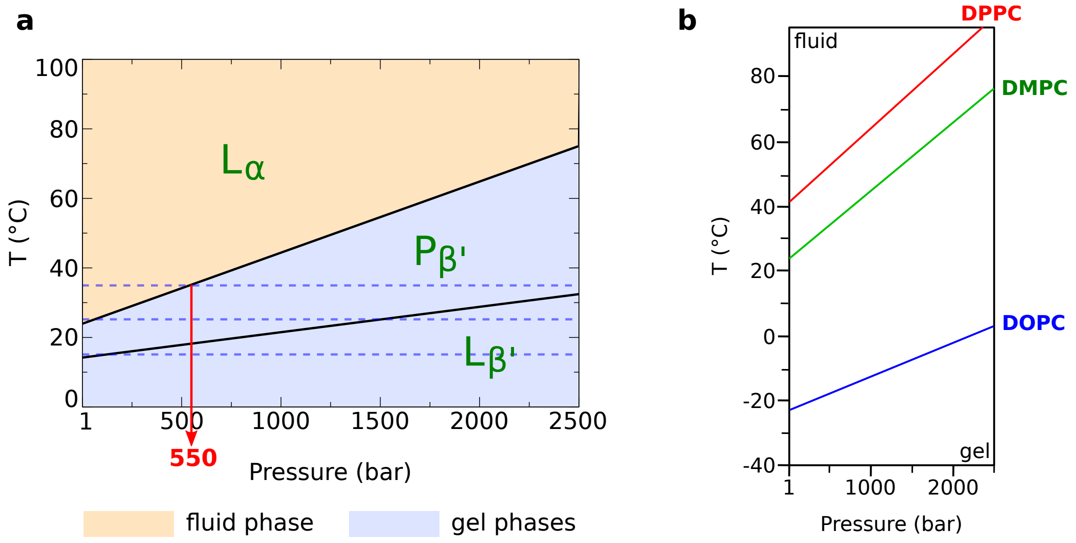


SUPPLEMENTARY FIG. 1. **Temperature/Pressure (T/P) phase properties of various lipid vesicles in excess water (in the absence of proteins)**. **a,** T/P phase diagram of hydrated multilamellar vesicles of DMPC. In the present study, the pressure ramps have been carried out along three isotherms (15, 25 and 35°C) represented by dashed blue lines. The fluid-like crystalline phase is referred to as $L_\alpha$ according to Luzzati's nomenclature[1]. It has a lamellar structure with conformationally disordered acyl chains. Additionally, two solid-like gel phases, $P_{\beta'}$ and $L_{\beta'}$, exist wherein the chains exhibit a more extended conformation. $P_{\beta'}$, commonly referred to as the ripple gel phase, is characterized by a periodic undulation of the bilayer within the lamellae plane. In this phase, the acyl chains are in a higher degree of ordering compared to the $L_\alpha$ phase, and they are tilted in relation to the normal to the bilayer plane. This phase occurs in bilayers consisting of phospholipids with saturated hydrocarbon chains[2], wherein they arrange themselves into a regular hexagonal lattice[3, 4]. $L_{\beta'}$ exhibits a lamellar bilayer structure characterized by fully extended and tilted hydrocarbon chains, but it is packed in a slightly distorted hexagonal lattice in comparison to $P_{\beta'}$[3, 4]. The gel-to-gel $P_{\beta'} \rightarrow L_{\beta'}$ pre-transition, which was predicted to happen at approximately 1500 bar at 25°C[5], was not observed in the present study possibly because the periodic length of the ripple phase is ∼145 Å[2, 6], i.e., larger than the diameter of the MSP1D1 nanodisc (∼10 nm). It is also possible that the similar relaxation properties in the $P_{\beta'}$ and $L_{\beta'}$ phases prevent any observable transition (this diagram is adapted from [5]). **b,** Fluid and gel phase demarcation lines for DPPC, DMPC and DOPC (based on an abundant literature). No such data is available for Δ9–cis–PC which is believed to remain in a fluid phase in the T/P conditions used in the present study using nanodiscs. This is due to the two cis-double bonds that introduce kinks and make it more difficult to align the fatty chains with each other reducing the ordering effect of pressure. Dipalmitoylphosphatidylcholine (DOPC), in which each acyl chain contains an additional 4 carbon atoms (18:1(Δ9–cis–PC)), has a $T_m$ = -23°C and ∼0°C at 1 and 2500 bar[7], respectively (blue line in the diagram). Based on these values, Δ9–cis–PC has necessarily lower $T_m$ due also to shorter fatty chains, by analogy with fully saturated DPPC (16 carbon atoms per acyl chain) (red line) and DMPC (green line) which have $T_m$ equal to 41 and 24°C at 1 bar, respectively.

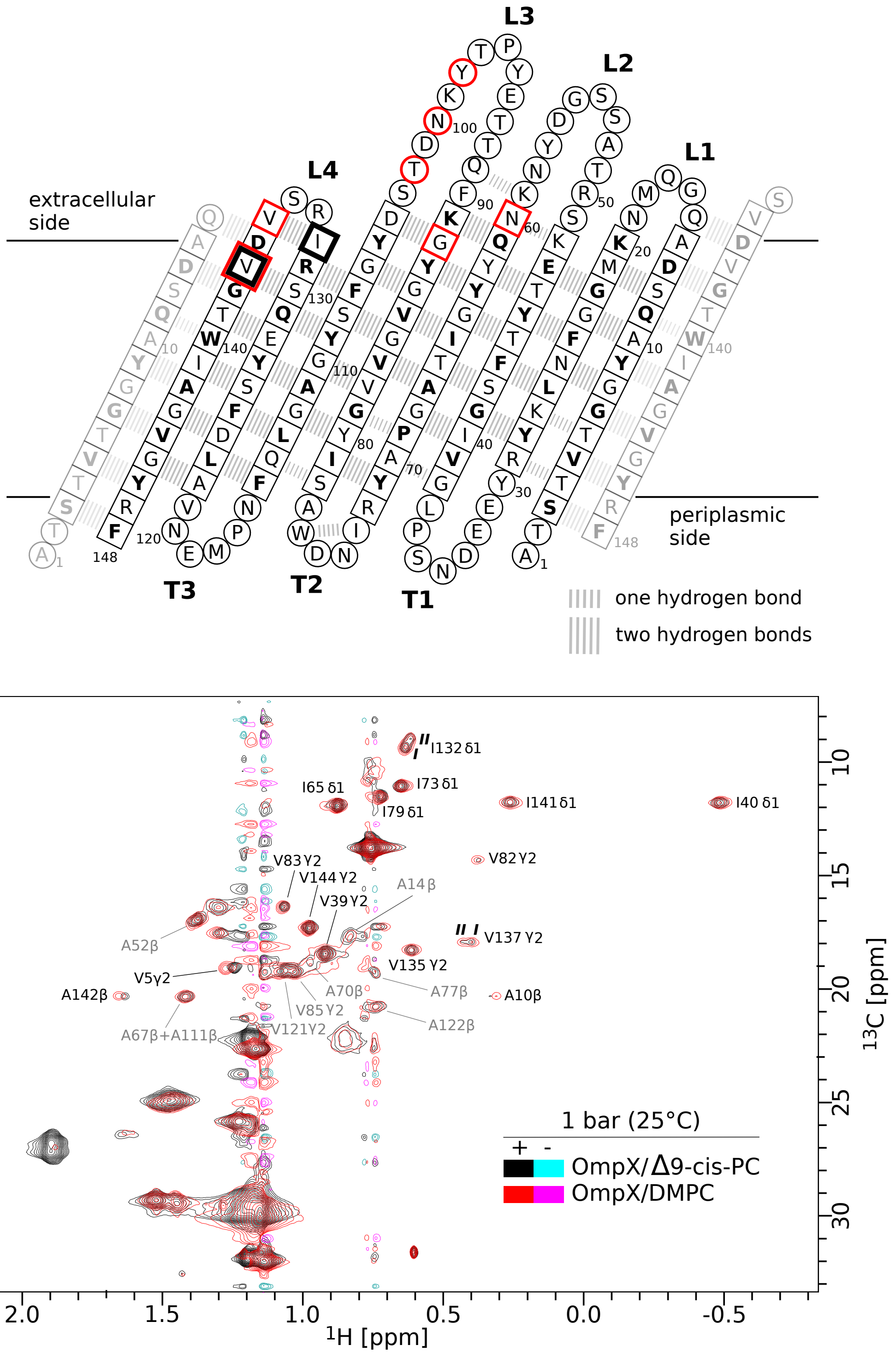


Supplementary Fig. 2. **NMR assignments of $^{13}CH_3$–$\beta$–Ala, $^{13}CH_3$–$\delta_1$–Ile and $^{13}CH_3$–$\gamma_2$(proS)–Val of OmpX in DMPC and $\Delta$9–cis–PC nanodiscs.** **Top**, Snake diagram of OmpX. Residues in $\beta$–strands are shown in squares, the others in circle. I132 and V137 that display splitted signals are framed with a bold square. Residues of the $\beta$-barrel that point their side chains to the lipid bilayer are written in bold black letters. The hydrogen bond network is based on OmpX NMR structure in MSP1D1 nanodiscs (PDB ID: 1QJ8[8]). **L** and **T** stands for extracellular loops and periplasmic turns, respectively. Residues involved in virulence and defense of the analog of OmpX protein Ail in *Y. enterocolitica* are squared/circled in red (from Vogt and Schulz[9]). **Bottom**, Two superimposed 2D $^{1}H$,$^{13}C$ SOFAST-HMQC[10] spectra collected at ambient pressure (1 bar) and 25°C. Residues labeled in grey were not taken into consideration due to either spectral crowding or lipid noise signals. The unlabeled signals correspond to lipid and lipoprotein $CH_n$ moieties as described in Supplementary Fig. 3.

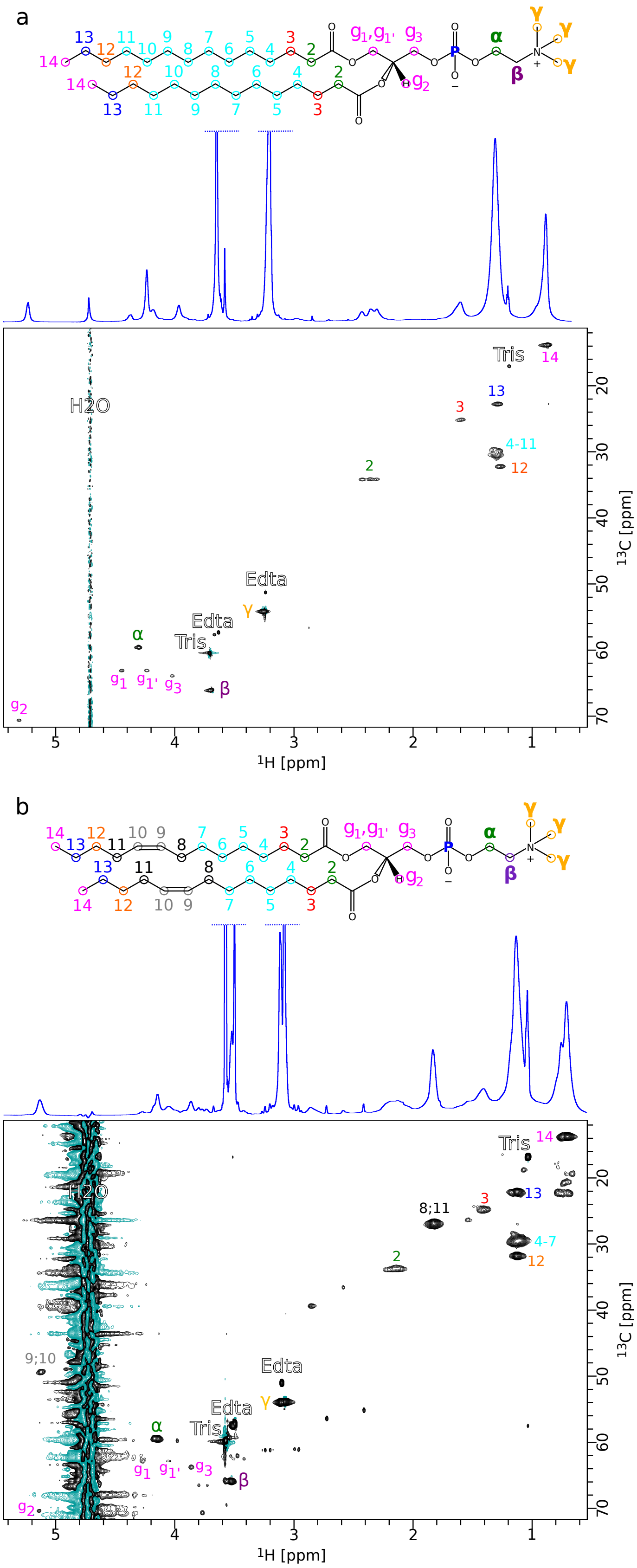


SUPPLEMENTARY FIG. 3. **$^1$H,$^{13}$C NMR assignments of DMPC (top) and Δ9–cis–PC (bottom) CH$_n$ in OmpX-devoid nanodiscs.** On top of each 2D $^1$H,$^{13}$C SOFAST-HMQC[10] spectrum is displayed a 1D $^1$H spectrum and the lipid chemical structure with a color code which is reproduced on the assignments. The assignments are based on 2D $^1$H,$^1$H TOCSY, NOESY and COSY experiments. The unlabeled signals correspond to the lipoprotein MSP1D1.

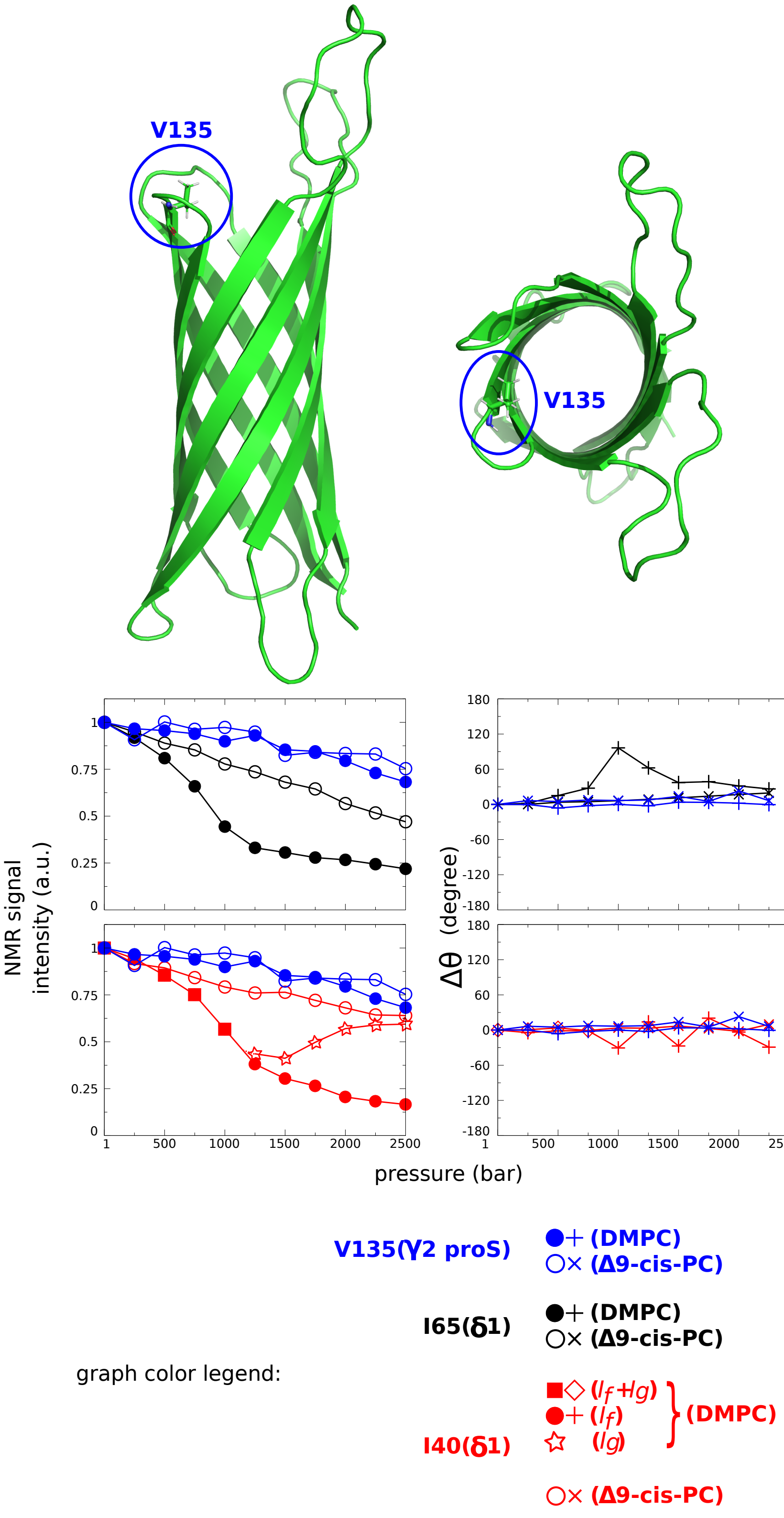


SUPPLEMENTARY FIG. 4. **Comparison of V135($\gamma_2$pro$S$) $^{13}CH_3$ barotropic behavior NMR signal with membrane and cavity oriented methyl groups.** V135 is located at the apex of the 8$^{th}$ strand and exposed to water. Among all the $^{13}CH_3$ of OmpX studied here, this methyl group represents an ideal reference as its NMR signal has been found to be the less impacted the lipid content, as shown by both the barotropic evolution of its intensity and pressure-dependence in $^{1}$H and $^{13}$C chemical shifts ($\Delta\theta$). **Top**, Location of V135 in OmpX. **Bottom**, Comparison of the barotropic evolution of $^{13}CH_3$ NMR intensities and chemical shifts ($\Delta\theta = \theta(P_i) - \theta(1\text{ bar})$; see the definition of $\theta$ in § 2) between V135($\gamma_2$pro$S$) and membrane-oriented I65($\delta_1$) and cavity-oriented I40($\delta_1$).

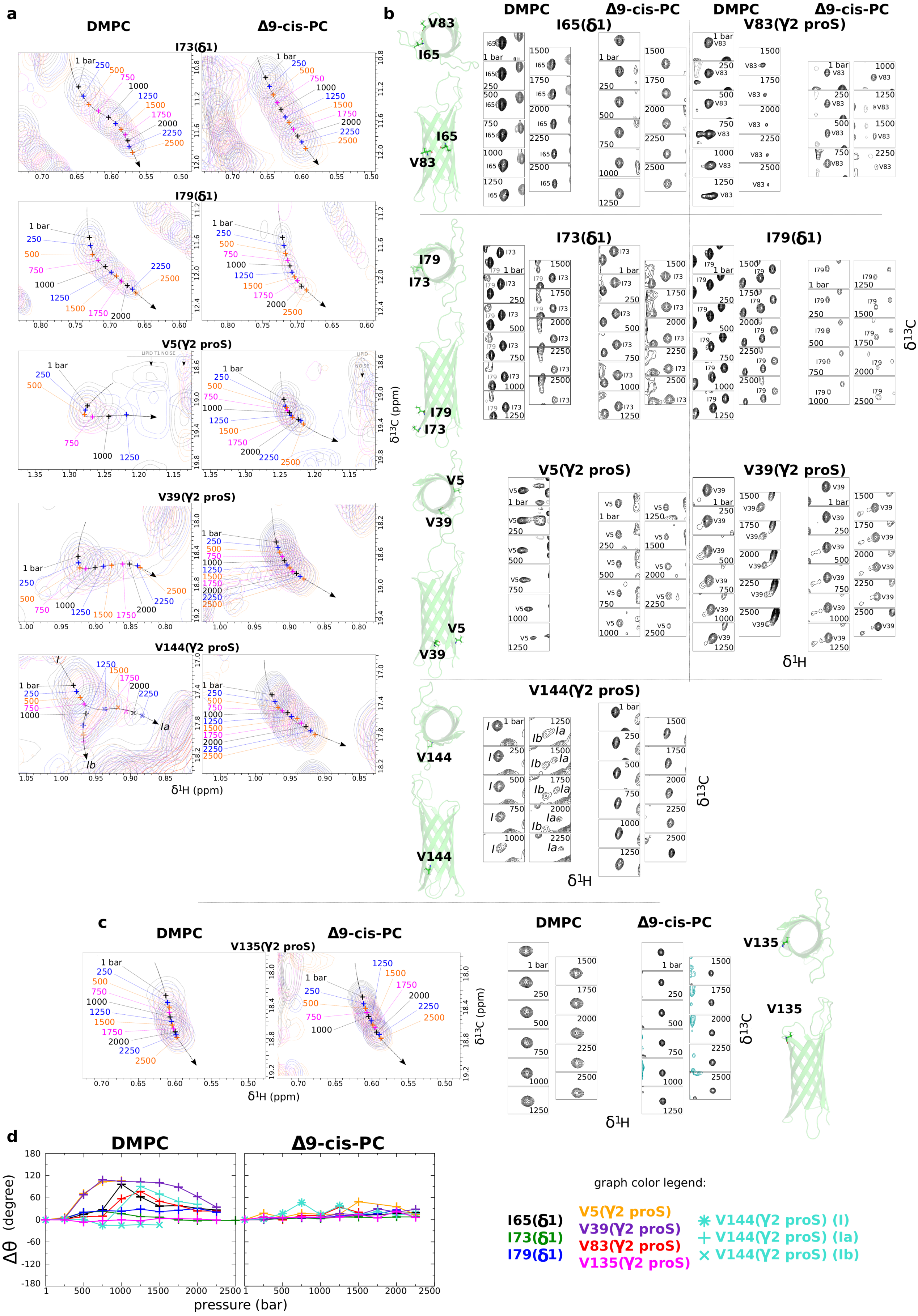
a
DMPC
Δ9-cis-PC
I73(δ1)
I79(δ1)
V5(γ2 proS)
V39(γ2 proS)
V144(γ2 proS)
δ1H (ppm)
δ13C (ppm)
b
I65(δ1)
V83(γ2 proS)
I73(δ1)
I79(δ1)
V5(γ2 proS)
V39(γ2 proS)
V144(γ2 proS)
δ1H
δ13C
c
V135(γ2 proS)
d
Δθ (degree)
pressure (bar)
graph color legend:
I65(δ1)
I73(δ1)
I79(δ1)
V5(γ2 proS)
V39(γ2 proS)
V83(γ2 proS)
V135(γ2 proS)
V144(γ2 proS) (I)
V144(γ2 proS) (Ia)
V144(γ2 proS) (Ib)

Supplementary Fig. 5. **Barotropic evolution of membrane-oriented $^{13}CH_3$ NMR signals of OmpX in DMPC and $\Delta$9–cis–PC nanometric bilayers at 25°C. a,** In complement to Fig. 4, Illustrations of superimposed 2D $^{1}H$,$^{13}C$ SOFAST-HMQC NMR spectra[10] for residues V5, V39, I73, I79 and V144. The numbers represent the hydrostatic pressures that were applied. All spectra have been represented at the same scale. For V144, *cf.* also Supplementary Fig. 12. **b,** Each panel represents successive 2D $^{1}H$,$^{13}C$ SOFAST-HMQC spectra acquired along the pressure ramp. They are all represented at the same scale, *i.e.*, spectral widths of 237.5 and 358.2 Hz, respectively in the $^{1}H$ and $^{13}C$ dimensions. For each amino acid, these spectra are represented superimposed to each other in **a** or Fig. 4a (for I65 and V83). **c,** Same as **a,b** for the reference residue V135. **d,** Representations of the barotropic evolution of the $^{13}CH_3$ chemical shift cross peaks through the angle $\theta$ ($\Delta\theta = \theta(P_i) - \theta(1\text{ bar})$; see the definition of $\theta$ in § 2).

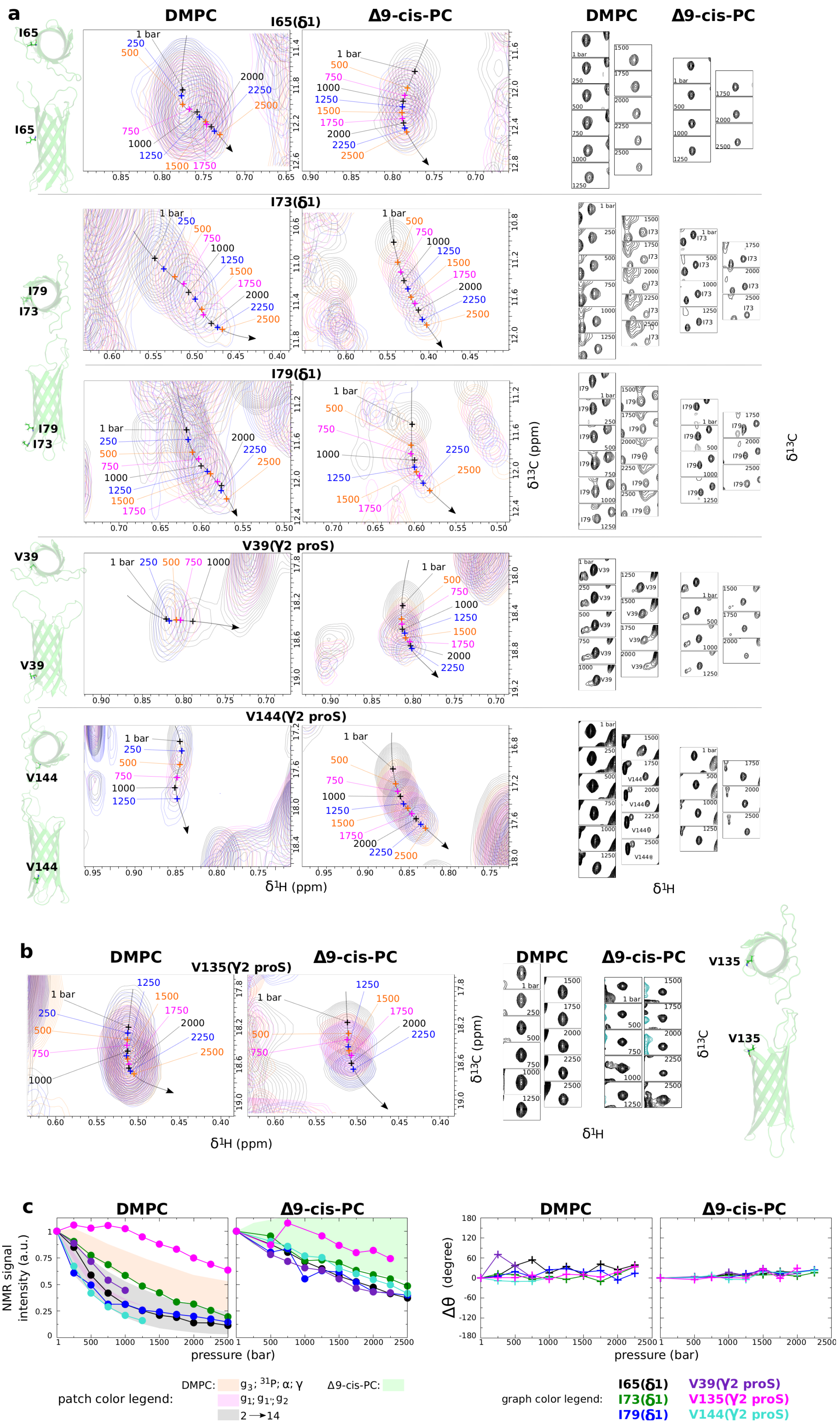
a
DMPC
Δ9-cis-PC
I65(δ1)
I73(δ1)
I79(δ1)
V39(γ2 proS)
V144(γ2 proS)
δ¹H (ppm)
δ¹³C (ppm)
b
V135(γ2 proS)
c
NMR signal intensity (a.u.)
pressure (bar)
Δθ (degree)
patch color legend:
DMPC: g3; ³¹P; α; γ
g1; g1'; g2
2 → 14
Δ9-cis-PC:
graph color legend:
I65(δ1)
I73(δ1)
I79(δ1)
V39(γ2 proS)
V135(γ2 proS)
V144(γ2 proS)

SUPPLEMENTARY FIG. 6. **Barotropic evolution of NMR signals of membrane-oriented $^{13}CH_3$ of OmpX in DMPC and $\Delta$9–cis–PC nanodiscs at 15°C. a** to **c**, Legend same as Supplementary Fig. 5 (V5 and V83 where not measured due to to low signal-to-noise ratio in the spectra at 15°C).

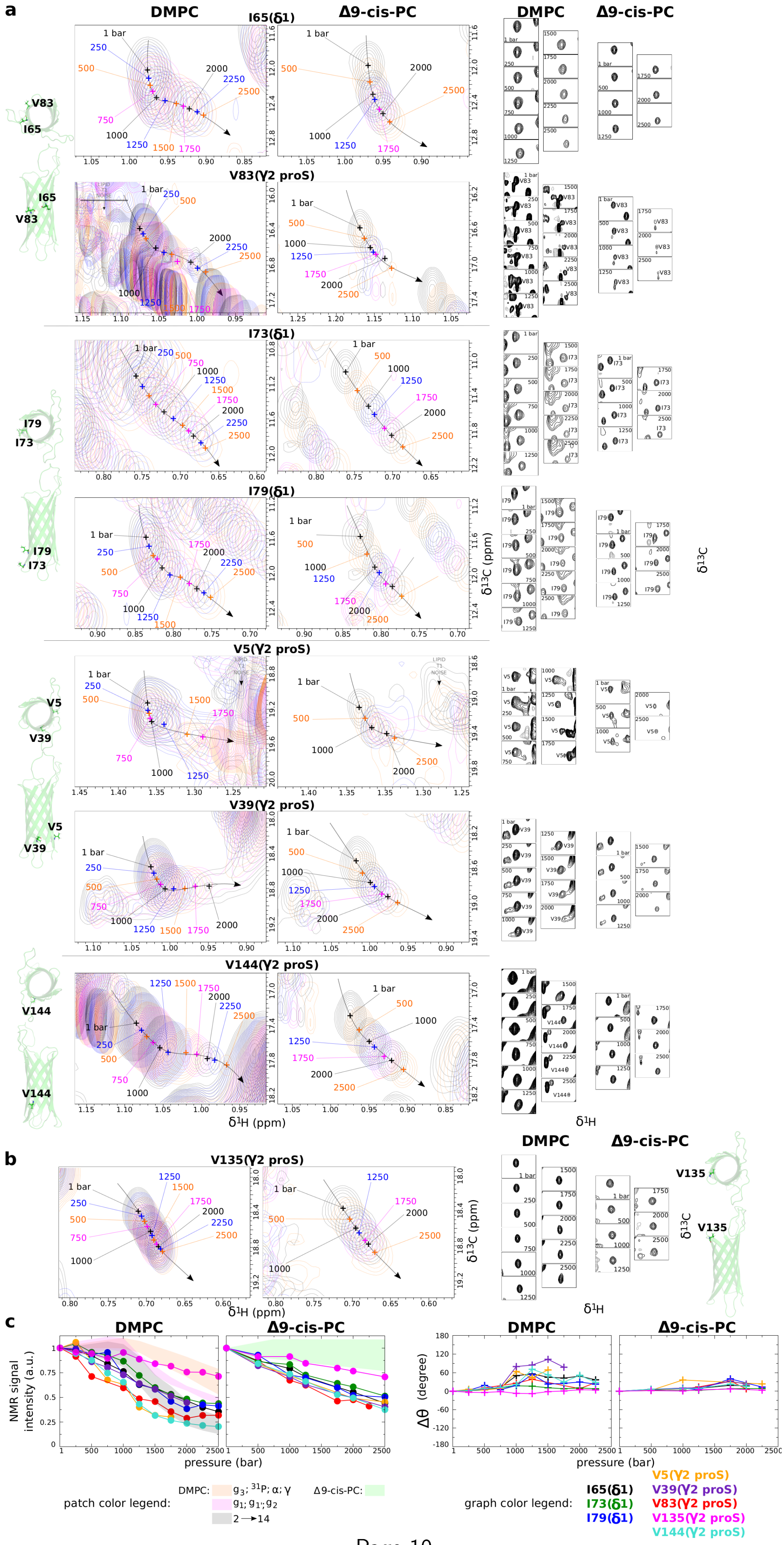
a
DMPC
Δ9-cis-PC
I65(δ1)
V83(γ2 proS)
I73(δ1)
I79(δ1)
V5(γ2 proS)
V39(γ2 proS)
V144(γ2 proS)
δ1H (ppm)
δ13C (ppm)
δ1H
δ13C
b
V135(γ2 proS)
c
NMR signal intensity (a.u.)
pressure (bar)
Δθ (degree)
patch color legend:
graph color legend:
I65(δ1)
I73(δ1)
I79(δ1)
V5(γ2 proS)
V39(γ2 proS)
V83(γ2 proS)
V135(γ2 proS)
V144(γ2 proS)

Supplementary Fig. 7. **Barotropic evolution of NMR signals of membrane-oriented $^{13}CH_3$ of OmpX in DMPC and △9–cis–PC nanodiscs at 35°C. a**, **b**, Legend same as Supplementary Fig. 5.

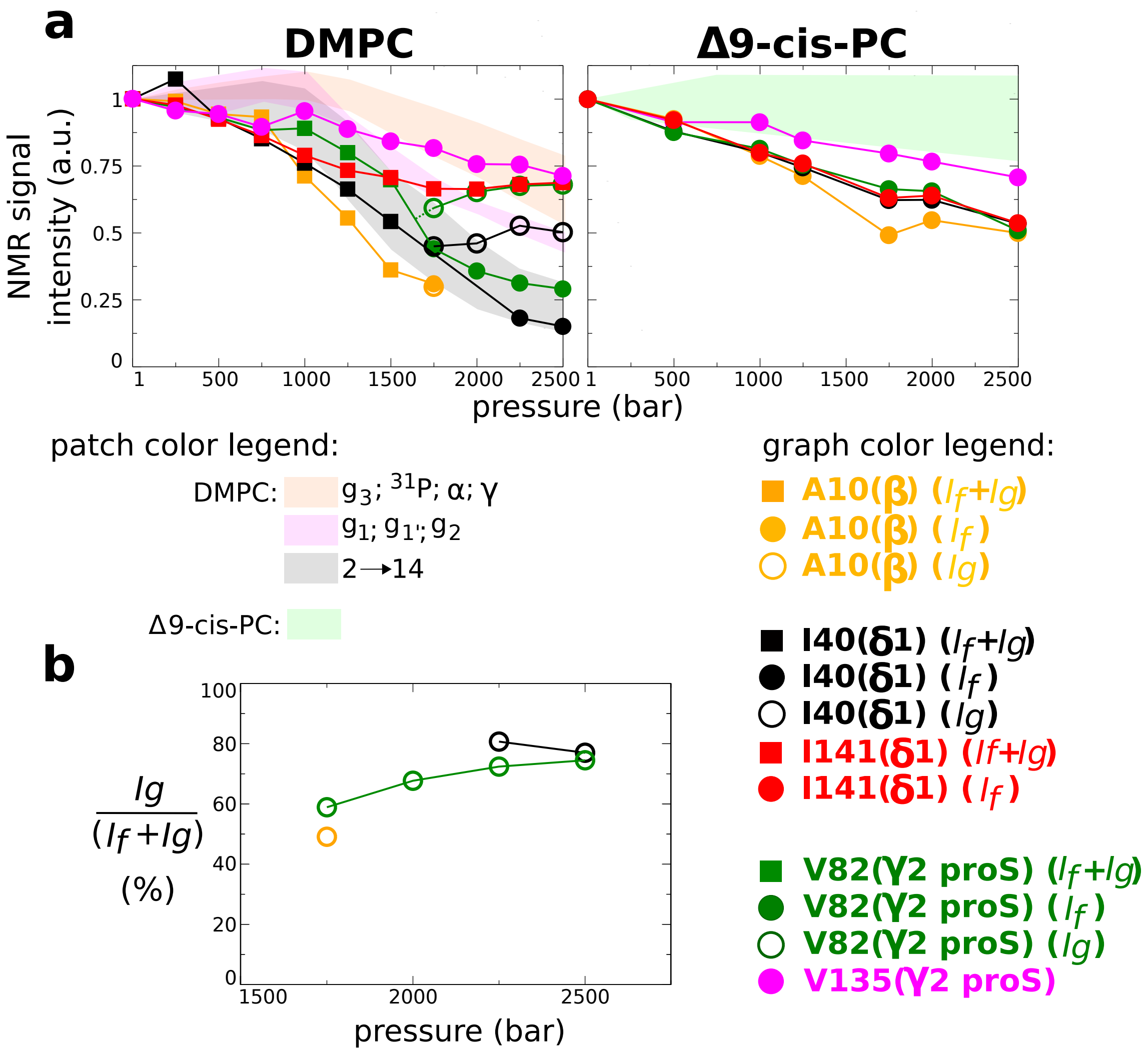


SUPPLEMENTARY FIG. 8. **Barotropic evolution of NMR signal intensities of barrel interior-oriented $^{13}CH_3$ of OmpX in DMPC and Δ9–cis–PC nanodiscs at 35°C (in complement to Extended Data Fig. 2).** **a**, Comparison of the barotropic evolutions of $^{13}CH_3$ NMR signal intensity. **b**, Barotropic evolution of the relative population of $I_g$.

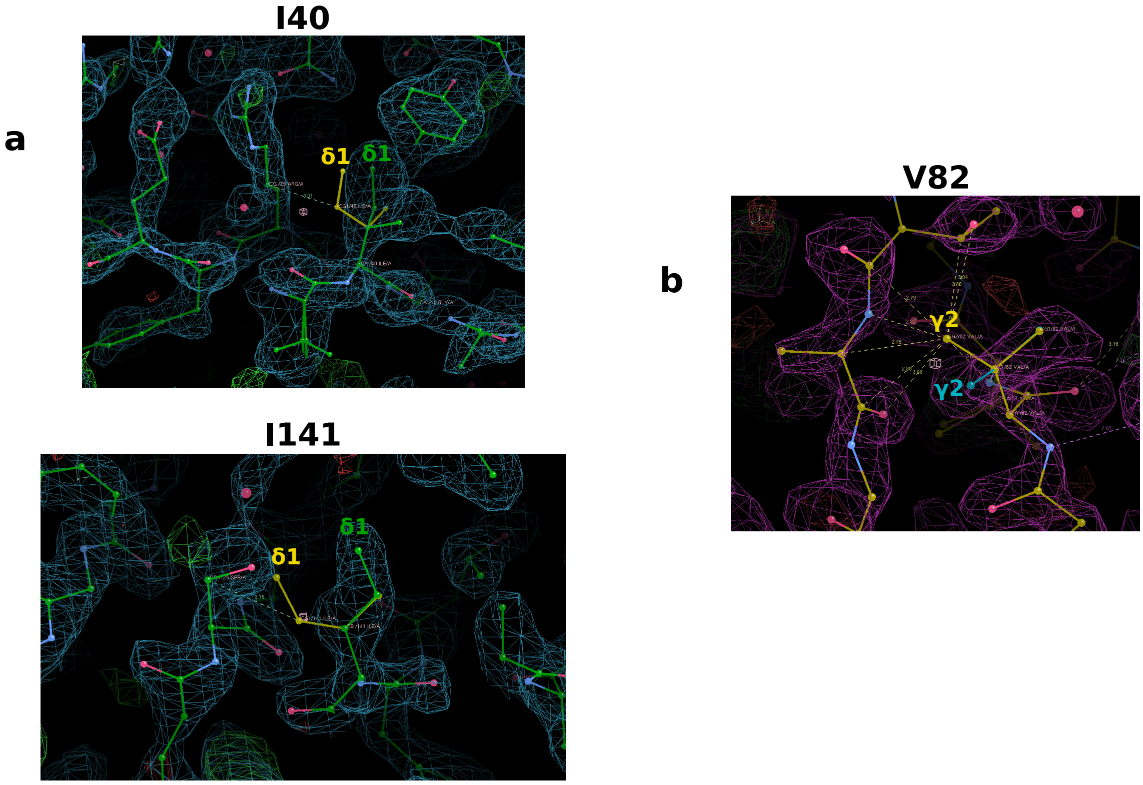


Supplementary Fig. 9. **Putative $\chi 1$ (N–C$\alpha$–C$\beta$–C$\gamma$) rotamers for cavity-membrane oriented I40, V82 and I141 side chains.** **a**, close-ups in the I40 (Top) and I141 (bottom) and V82 (**b**) regions. Amino acids are represented in a ball and stick representation. The electron density is represented in *blue* (**a**) or *purple* (**b**) from OmpX crystal structure (PDB ID: 1QJ8). In *green* (**a**) or *blue* (**a**,**b**), the amino acids correspond to the original structure and in *yellow* (**a** and **b**) is represented for each panel one alternative $\chi 1$ rotamer causing no additional clash. A rotamer with a side chain displaying $^{13}$C–$^{13}$C distances $\geq$ 3Å with its neighborhood is considered conceivable. In the case of I141, there is a steric clash with S126 that would require a reorientation of S126 side chain. Representations and rotamer calculation have been generated by *Coot* software[11].

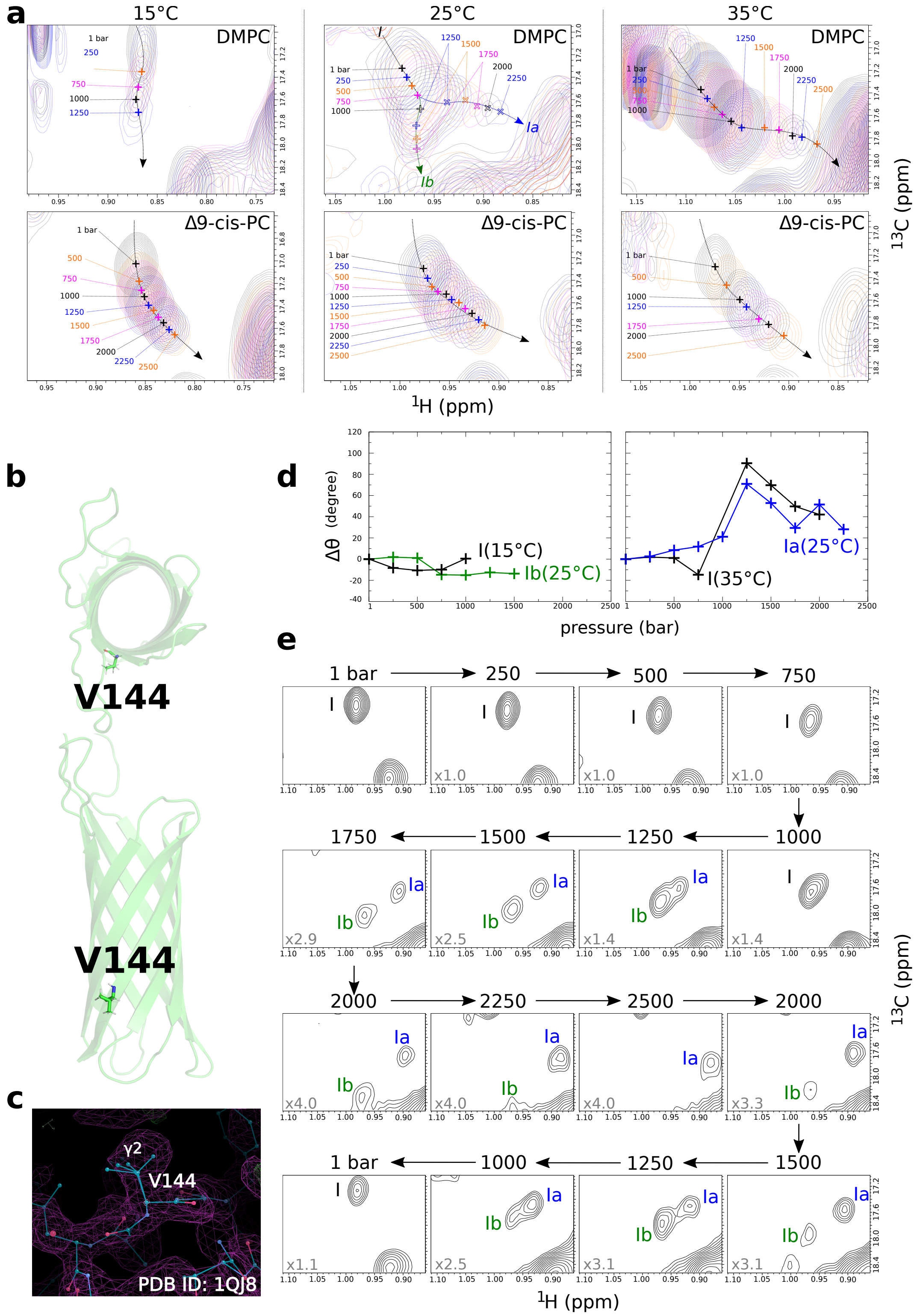


SUPPLEMENTARY FIG. 10. **Comparison of the barotropic evolution of $\gamma_2$-pro$S$-V144 methyl group observed at 15, 25 and 35°C. a**, Superimposed 2D $^{1}$H,$^{13}$C SOFAST-HMQC NMR spectra. The numbers represent the hydrostatic pressures that were applied. All spectra have been represented at the same EL: intensity? scale. **b**, Position of V144 denoted on cartoon representations of OmpX, observed from both a parallel and perpendicular axis (from the extracellular side) to the plane of the membrane. **c**, Two possible orientations of V144 isopropyl group in OmpX crystal structure represented with balls and sticks in the electron density (PDB ID: 1QJ8). **d**, Comparison of the pressure-dependence of methyl combined $^{1}$H and $^{13}$C NMR chemical shifts, $\Delta\theta$, between $I$(15°C) and $I_b$(25°C) signals (left) and between $I$(35°C) and $I_a$(25°C) (right) (se $\theta$ definition in § 2). **e**, Sequential description of the reversible barotropic evolution of $I$, $I_a$ and $I_b$ signals at 25°C.

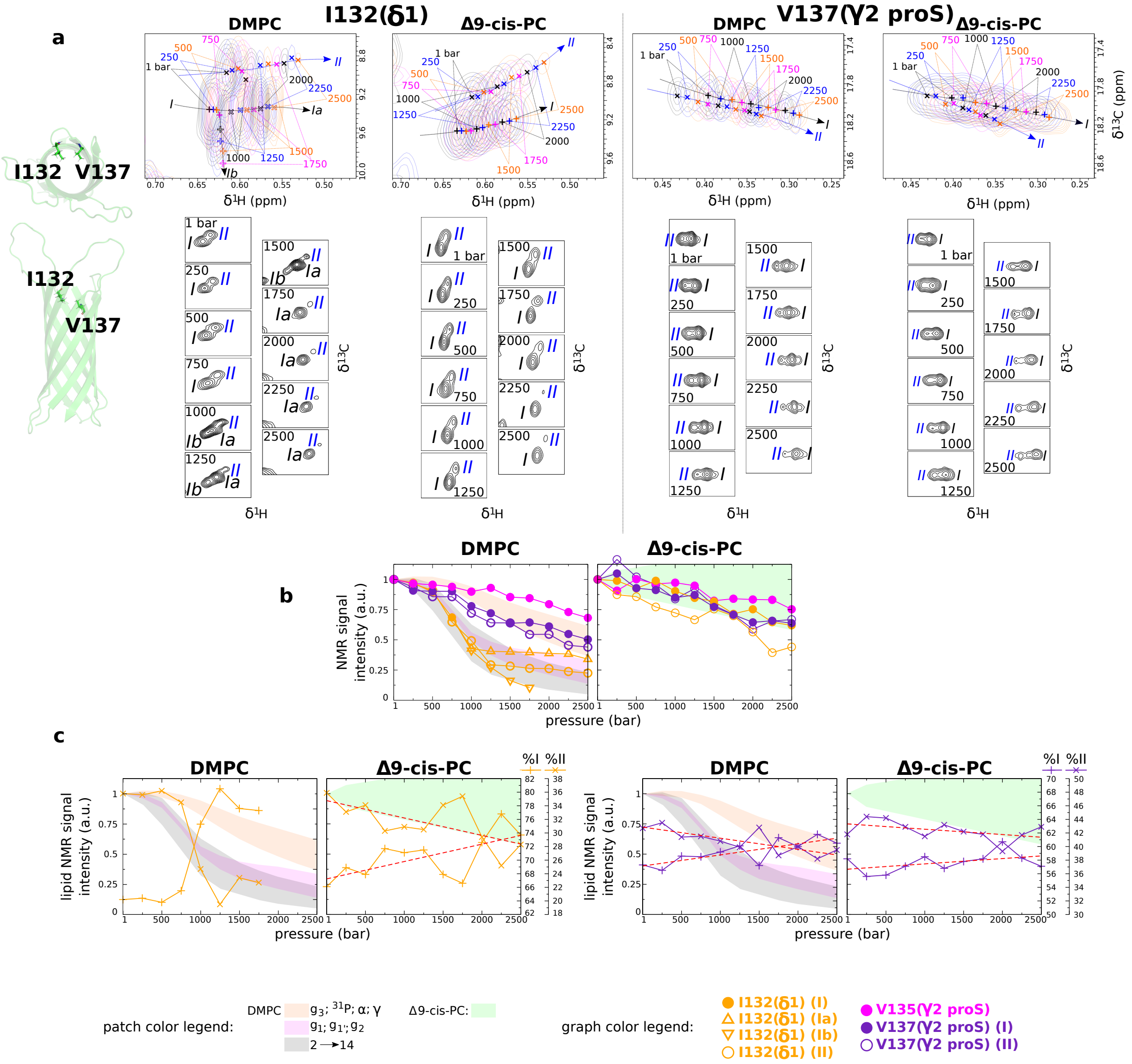


SUPPLEMENTARY FIG. 11. **Barotropic evolution of I132($\delta_1$) and V137($\gamma_2$proS) $^{13}CH_3$ NMR signals of OmpX in DMPC and Δ9–cis–PC nanometric bilayers at 25°C. a,** Superimposed 2D $^1$H,$^{13}$C SOFAST-HMQC NMR spectra[10] (**Top**) and successive 2D $^1$H,$^{13}$C SOFAST-HMQC spectra acquired along the pressure ramp (**Bottom**). All spectra are represented at the same scale, *i.e.*, spectral widths of 237.5 and 358.2 Hz, respectively in the $^1$H and $^{13}$C dimensions. Numbers indicate the pressure applied. **b,** Evolution of NMR signal intensities under pressurization. **c,** Evolutionary trajectories of populations *I* and *II* for I132 (left) and V137 (right). The red dashed lines represent linear regression fits.

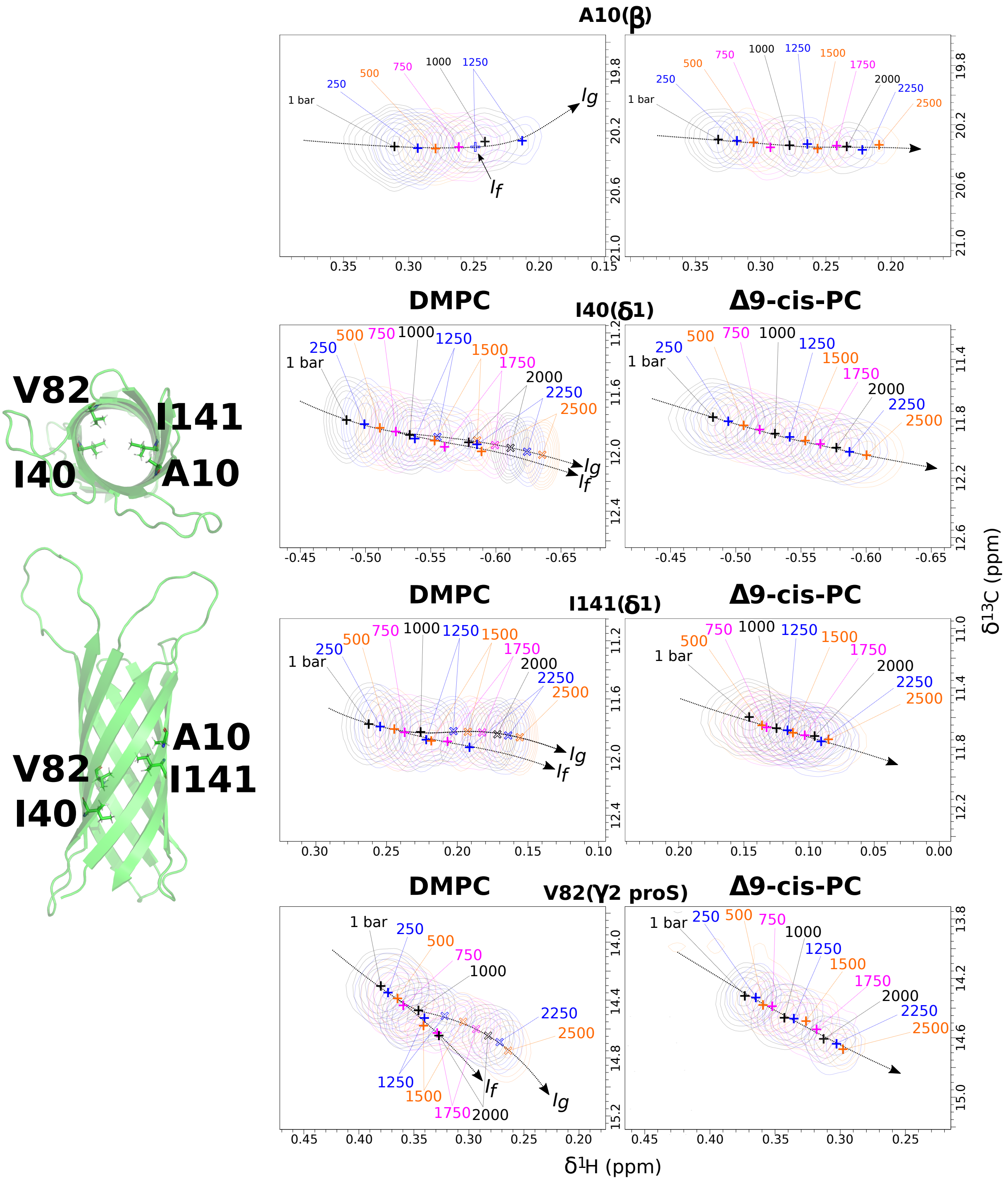


Supplementary Fig. 12. **Barotropic evolution of interior cavity-oriented $^{13}CH_3$ NMR signals of OmpX in DMPC and Δ9–cis–PC nanometric bilayers at 25°C (in complement to Fig. 5).** Illustrations of superimposed 2D $^{1}H$,$^{13}C$ SOFAST-HMQC NMR spectra[10] for residues A10, I40, V82 and I141. The numbers represent the hydrostatic pressures that were applied. All spectra have been represented at the same scale. $I_f$ and $I_g$ refer to fluid and gel states, respectively (see also data concerning the reference residue V135 in Supplementary Fig. 5c).

#### 2 Analysis of pressure-induced evolution of OmpX $^{13}CH_3$ chemical shifts: definition of the angle $\theta$

To visualize the impact of pressure on the evolution of $^{13}CH_3$ chemical shifts, the angle $\theta$ is defined as follows:

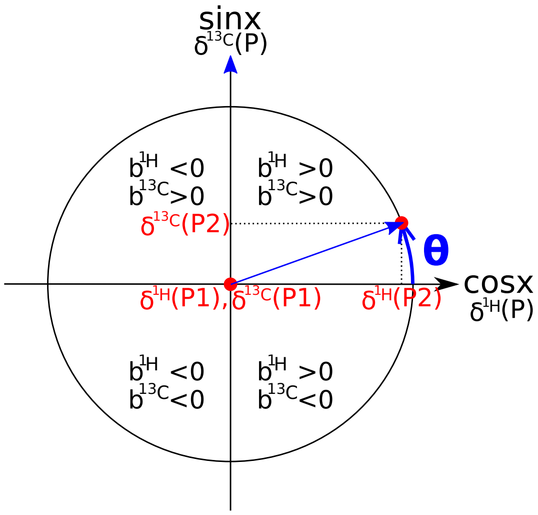


$$cos\theta = \frac{\delta^{1H}(P2)-\delta^{1H}(P1)}{[(\delta^{1H}(P2)-\delta^{1H}(P1))^2+[(\delta^{13C}(P2)-\delta^{13C}(P1))\times\frac{\gamma^{13C}}{\gamma^{1H}}]^2]^{1/2}}$$

$$sin\theta = \frac{[\delta^{13C}(P2)-\delta^{13C}(P1)]\times\frac{\gamma^{13C}}{\gamma^{1H}}}{[(\delta^{1H}(P2)-\delta^{1H}(P1))^2+[(\delta^{13C}(P2)-\delta^{13C}(P1))\times\frac{\gamma^{13C}}{\gamma^{1H}}]^2]^{1/2}}$$

where, $b^{1H} = \frac{\delta^{1H}(P2)-\delta^{1H}(P1)}{\Delta P}$ ; $b^{13C} = \frac{\delta^{13C}(P2)-\delta^{13C}(P1)}{\Delta P}$

For a given pressure range ($\equiv P2-P1$), $\theta$ contains information on the linear coefficient $b$ of both the proton ($b^{1H}$) and $^{13}$C ($b^{13C}$). As a first approximation, we consider the *b* coefficients as an average value in the pressure range $P2-P1$. In our case, $P2-P1$ is comprised between 100 and 250 bar.

so,

$$cos\theta = \frac{b^{1H}}{[(b^{1H})^2+(b^{13C}\times\frac{\gamma^{13C}}{\gamma^{1H}})^2]^{1/2}} \text{ or } \frac{b^{1H}}{[(b^{1H})^2+(b^{15N}\times\frac{\gamma^{15N}}{\gamma^{1H}})^2]^{1/2}}$$

$$sin\theta = \frac{b^{13C}\times\frac{\gamma^{13C}}{\gamma^{1H}}}{[(b^{1H})^2+(b^{13C}\times\frac{\gamma^{13C}}{\gamma^{1H}})^2]^{1/2}} \text{ or } \frac{b^{15N}\times\frac{\gamma^{15N}}{\gamma^{1H}}}{[(b^{1H})^2+(b^{15N}\times\frac{\gamma^{15N}}{\gamma^{1H}})^2]^{1/2}}$$

with:

$\gamma^{1H}$ = 267.52218744$\times10^{-6}$ rad$\times$s$^{-1}\times$T$^{-1}$ and $\gamma^{13C}$ = 67.2828$\times10^{-6}$ rad$\times$s$^{-1}\times$T$^{-1}$

⋆ if cos$\theta$ >0 (b$^{1H}$ >0) and sin$\theta$ >0 (b$^{13C}$ >0):

$$\theta^{13CH_3} = arccos(\frac{b^{1H}}{[(b^{1H})^2+(b^{13C}\times\frac{\gamma^{13C}}{\gamma^{1H}})^2]^{1/2}}) \equiv arcsin(\frac{b^{13C}\times\frac{\gamma^{13C}}{\gamma^{1H}}}{[(b^{1H})^2+(b^{13C}\times\frac{\gamma^{13C}}{\gamma^{1H}})^2]^{1/2}})$$

⋆ if cos$\theta$ <0 (b$^{1H}$ <0) and sin$\theta$ >0 (b$^{13C}$ >0):

$$\theta^{^{13}CH_3} = arccos(\frac{b^{^1H}}{[(b^{^1H})^2+(b^{^{13}C}\times\frac{\gamma^{13C}}{\gamma^{1H}})^2]^{1/2}}) \equiv \Pi - arcsin(\frac{b^{^{13}C}\times\frac{\gamma^{13C}}{\gamma^{1H}}}{[(b^{^1H})^2+(b^{^{13}C}\times\frac{\gamma^{13C}}{\gamma^{1H}})^2]^{1/2}})$$

⋆ if cos$\theta$ <0 (b$^{^1H}$ <0) and sin$\theta$ <0 (b$^{^{13}C}$ <0):

$$\theta^{^{13}CH_3} = -arccos(\frac{b^{^1H}}{[(b^{^1H})^2+(b^{^{13}C}\times\frac{\gamma^{13C}}{\gamma^{1H}})^2]^{1/2}}) \equiv -\Pi - arcsin(\frac{b^{^{13}C}\times\frac{\gamma^{13C}}{\gamma^{1H}}}{[(b^{^1H})^2+(b^{^{13}C}\times\frac{\gamma^{13C}}{\gamma^{1H}})^2]^{1/2}})$$

⋆ if cos$\theta$ >0 (b$^{^1H}$ >0) and sin$\theta$ <0 (b$^{^{13}C}$ <0):

$$\theta^{^{13}CH_3} = -arccos(\frac{b^{^1H}}{[(b^{^1H})^2+(b^{^{13}C}\times\frac{\gamma^{13C}}{\gamma^{1H}})^2]^{1/2}}) \equiv arcsin(\frac{b^{^{13}C}\times\frac{\gamma^{13C}}{\gamma^{1H}}}{[(b^{^1H})^2+(b^{^{13}C}\times\frac{\gamma^{13C}}{\gamma^{1H}})^2]^{1/2}})$$

To facilitate the comparison of $\theta$ between different $^{13}$CH$_3$, we define $\Delta\theta$ in order to have a $\Delta\theta$ equal to 0 at ambient pressure for all methyls:

$$\Delta\theta = \theta(P_i) - \theta(1\ \text{bar})$$
